# Glucose and fructose differently mediate alcohol cocktail drinking in female and male rats: interaction of glucose and alcohol on post-ingestive behavior

**DOI:** 10.64898/2026.04.15.718765

**Authors:** Isabel R. K. Kuebler, Gerin Zimmerman, Shi Q. Ng, Hannah M. Schneider, Kyle Sextro, Audrey Denning, Bryn Mattes, Matthew Matuszeski, Mauricio Suarez, Ken T. Wakabayashi

## Abstract

Sweetened alcoholic beverages are thought to contribute to developing Alcohol Use Disorder by increasing palatability. One monosaccharide, glucose, readily enters the brain more than fructose and directly impacts the activity of central neurons. The objective of this study is to determine the impact of glucose versus fructose on alcohol drinking patterns in female and male rats. Rats drank alcohol cocktails (1.25%-10%) containing either glucose or fructose (10%) in 4-hour sessions. We sought to parse orosensory effects from post-ingestive central effects by analyzing drinking microstructure. We compared measures of palatability and post-ingestive feedback between early and later in the session when brain levels of alcohol and glucose are different. We found that rats of both sexes drank more low alcohol glucose cocktails than cocktails containing fructose by volume and by overall calories. When considering the dose of alcohol, glucose potentiated alcohol intake by shifting the dose-response curve leftward compared to similar fructose cocktails. We found that drinking patterns associated with palatability remained stable for both types of cocktails over the entire drinking session. In contrast, post-ingestive behavior related to brain mediated satiety or positive feedback showed a greater influence of the session time, as well as a greater interaction with sex. Overall, our results suggest that glucose and alcohol interact to impact central regulation of cocktail drinking. This highlights that the type of sugar within cocktails interacts and ultimately have different effects on brain regulated alcohol drinking.

## Introduction

In the United States, about 29.5 million people, or 10.5% of the population, aged 12 and older meet diagnostic criteria for alcohol use disorder (AUD) (SAMHSA, 2023). This statistic includes 753,000 adolescents aged 12-17 (SAMHSA, 2023). More broadly, alcohol drinking is prevalent among American high school students, 22.1% of whom report having at least one drink of alcohol in the last month (CDC, 2024). Within this sample, more girls and women (24.4%) report drinking than boys and men (20.0%) (CDC, 2024). Importantly, alcohol drinking that increases the risk of developing AUD often starts in early adolescence, with 13.3% of high school students reporting that their first drink of alcohol was before 13 years of age (CDC, 2024). Moreover, 8.8% of high schoolers report engaging in episodic drinking within the last month, with again girls and women reporting greater episodic drinking (10.0%) than boys and men (7.7%) (CDC, 2024); frequent episodic drinking in early adulthood (17-25 years of age) has been associated with higher rates of AUD, a decade later (Sloan et al., 2011), as well as in midlife (Patrick et al., 2023).

Although there are many social factors contributing to alcohol consumption and the development of AUD, one important consideration is that a wide range of alcoholic beverages contain additional sugar-based sweeteners. For example, flavored alcoholic beverages (FABs), commonly referred to as “ready-to-drink cocktails” or “alcopops,” are a loosely defined range of drinks that vary widely in their alcohol concentration and additive sweeteners. FABs are often studied in the context of adolescence because of the prevalence of sweetened FAB consumption in this age group (Griffiths and Sutherland, 1998; Fortunato et al., 2014; Albers et al., 2015). The assumption is that adolescent preference towards sweetened FABs is related to initiation and acquisition of alcohol drinking, mirroring preclinical research (Samson et al., 1989). Indeed, the role of added sweeteners like sugars in FABs in influencing alcohol drinking in both humans and preclinical models is explained partially in terms of taste and palatability. However, the reasons why any age group drinks FABs appear to be more complex (Jones and Reis, 2012; IRI, 2022). Further, focusing only on taste as the main component of FAB consumption diminishes the differing contribution of sugars used in FABs towards direct effects in the brain.

Different FABs commonly contain glucose and fructose monosaccharides in varying proportions, ranging from 42%-70% fructose versus glucose, with the most prevalent mixture in commercial beverages being 55% fructose, 45% glucose (FDA, 2018). Moreover, in many animal studies, the disaccharide sucrose is used in well-established “sweetened fade” procedures to facilitate acquisition of alcohol consumption and self-administration of unadulterated alcohol (Samson, 1986). Sucrose is, at times, considered equivalent to a 50% /50% glucose-fructose formulation, perhaps because of the monosaccharide constituents of sucrose as well as their caloric value. However, in solution, the glucose and fructose monomers making up sucrose remain bound together until they are broken down by gastric activity or enzymes in the small intestine. Consequently, glucose and fructose drank in commercial FABs and sucrose used in preclinical research ought to be considered different compounds until they enter the bloodstream. Together, differences in palatability across sugar type as well as the varying amounts of fructose versus glucose in beverages indicate that glucose and fructose should be studied separately from sucrose.

This consideration is critical because glucose and fructose differ in their central pharmacokinetics and pharmacodynamics. Transport of glucose and fructose through the blood-brain barrier differ by transporter type and speed, so that glucose arrives in the brain much faster and in much larger amounts than fructose (Duelli and Kuschinsky, 2001). Once within the brain, glucose is known to act directly on several biochemically and functionally distinct populations of neurons implicated in AUD (Burdakov et al., 2005; Cippitelli et al., 2010; Karlsson et al., 2016). Direct pharmacodynamic effects of fructose are unknown. Moreover, glucose and fructose drinking differ behaviorally: male rats escalate glucose drinking more than fructose, and both male and female rats develop a preference for glucose over fructose (Ackroff and Sclafani, 1991; Wakabayashi et al., 2016).

While sugars are well-established in facilitating alcohol drinking in rats, (Ayoub et al., 2020), separate examination of the effects of glucose and fructose on alcohol drinking has been rare. It has been recently reported that addition of a 25% solution of 55% fructose/42% glucose high fructose corn syrup potentiates intraoral operant self-administration in adult male Sprague-Dawley rats (Ayoub et al., 2020). However, the impact of glucose or fructose by themselves on alcohol drinking is less clear.

Glucose, fructose, and alcohol drinking in rats has traditionally been studied with an emphasis on orosensory taste, hedonic palatability, and licking pattern microstructure. Some measures have established that in male Sprague Dawley rats, fructose is generally more palatable than glucose (Davis, 1973), while another group has reported the opposite (Schier and Spector, 2016). Alcohol drinking using the same microstructure elements is reported to be comparatively aversive (Bice et al., 1992; Lin et al., 2013). Importantly, most of the data on drinking microstructure of these compounds is limited to short sessions (30-90 minutes), as they are structured to limit the post-ingestive contributions to drinking behavior. However, in studies focused on preclinical models of AUD-like behavior where the focus is on the central, post-ingestive effects of alcohol, longer multi-hour sessions are more common (Gilbert, 1974; Gill et al., 1986; Wolffgramm, 1990).

Therefore, the objective of the present study is to determine the impact of glucose and fructose on concentration-dependent alcohol drinking and drinking microstructure in Wistar rats. We chose to systematically compare the drinking behavior of both female and male rats during a longer multi-hour session more commonly used to model AUD-like behavior. This approach allowed us to parse orosensory mechanisms from post-ingestive effects, including central effects, that we hypothesized would be important in regulating alcohol-sugar cocktail drinking behavior.

## Material and methods

### Subjects

Data from 32 male and 32 female Wistar rats (Envigo), age matched (∼2.5-3 months) weighing 394 ± 23 g (male) and 289 ± 12 g (female) at the beginning of testing were used in this study. Female rats were intact and freely cycling. Because we had no *a priori* hypotheses on an interaction between ovarian cycle phase and the experimental variables, and there is currently no data on sex differences with sugar-sweetened alcohol drinking, we did not control the cycle phase. Rats were group housed upon arrival in a climate-controlled animal colony in filtered air cages and maintained on a 12-12 light-dark cycle (lights on at 15:00). Food and water were available *ad libitum* in the home cage. All procedures were approved by the University of Nebraska-Lincoln Animal Care and Use Committee and complied with the Guide for the Care and Use of Laboratory Animals (National Research Council (U.S.) et al., 2011). The ARRIVE Guidelines 2.0 were also considered during the submission of this manuscript. Sample size was determined after completing an initial pilot study and conducting a power analysis using 80% power and alpha=0.05 (G*Power version 3.1.9.7).

### Stereotaxic surgery

Rats were infused with an adenovirus associated vector (AAV) for future activation of lateral hypothalamic melanin-concentrating hormone neurons (bilaterally 0.5 µL, AP: -2.9 mm, ML +/-1.6 mm, DV -8.8 mm) using a Designer Receptor Exclusively Activated by Designer Drugs (DREADD) (20 male, 22 female), or DREADD-free controls (12 male, 10 female) in a procedure described elsewhere (Wakabayashi et al., 2019; Shields et al., 2021). Data from these DREADD manipulations were not analyzed in this manuscript; however, we mention this surgery here for rigor and replicability. Rats were allowed a minimum of 5 days postoperative recovery before drinking procedures (**Fig. 1a**).

**Figure 1.**
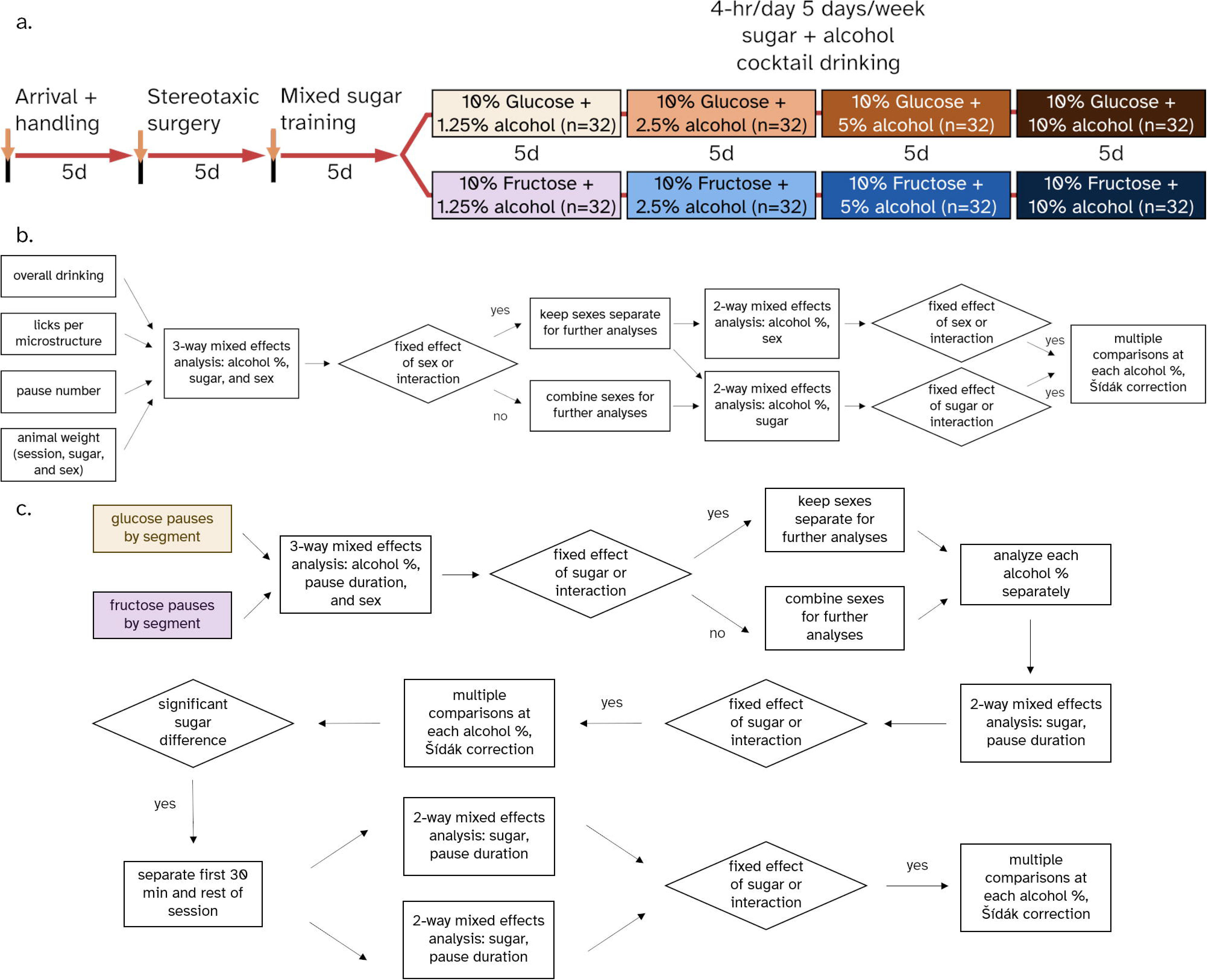
Methods. **(a)** Timeline of experimental procedures. **(b)** Statistical strategy used for all analyses including all pause lengths. Each time interval within the session was analyzed separately using this workflow for microstructure analyses. **(c)** Statistical strategy used to analyze number of pauses at various pause lengths

### Apparatus – alcohol intake chambers

Rat testing chambers (Med-Associates) were housed in sound-attenuating cubicles and equipped with two infrared lickometers on one wall of the chamber. A dim white houselight was located on the opposite wall. During all sessions, a single drinking bottle (∼250 mL) was mounted in the infrared lickometer furthest away from the door, with the lickometer spout 7.6 cm above the grid floor.

### Solutions

All alcohol cocktail solutions were prepared every 2-3 days with filtered food-grade water. Solutions used during this study were: a 5% w/v fructose (Sigma-Aldrich) and 5% glucose solution (Oakwood) (0.28M, respectively), a 10% w/v fructose or 10% w/v glucose solution (0.56 M) containing 1.25%, 2.5%, 5%, or 10% (v/v) alcohol, diluted from 190 proof ethyl alcohol, (ACS/USP grade, Decon Labs, Inc.).

### Shaping and intake sessions

Sessions were from 9 am-1 pm or 1:30-5:30 pm in two cohorts per day, 5 days per week (**Fig. 1a**). Prior to sessions analyzed for this study, rats were habituated to the intake chambers and the drinking procedure with a 5% fructose + 5% glucose solution for 5 days, or until all rats had acquired robust drinking behavior. Then, rats in each cohort were equally split into separate fructose cocktail and glucose cocktail groups for the rest of the intake sessions based on sugar drinking. Next, rats drank 5% fructose or 5% glucose solutions with first 1.25% alcohol the first week, then increasing to 2.5%, 5%, and 10% alcohol each of the following weeks.

### Data Processing, Management and Statistical Analysis

Individual licks during the session were recorded at a 10 ms temporal resolution. After the behavioral sessions, lickometer timestamp data were initially processed using a custom Konstanz Information Miner (KNIME) (Berthold et al., 2007) script adapted from a previously published R script (Raymond et al., 2018). Based on pilot data, licks with inter-lick intervals (ILIs) shorter than 90 ms were labeled as artifacts (i.e. “double licks”) due to the configuration of the lickometer apparatus. These licks were removed, and the ILI was recalculated using the next lick 90 ms or more after the first lick. Licks were then classified into two different microstructure classifications based on ILI length. These were “***clusters***” (groups of licks separated by an ILI > 500 ms, Davis, 1989), and “***runs***” (groups of licks separated by an ILI > 1s, Davis, 1989). We examined the shorter microstructure length, “***clusters***”, as they are lengths most typically used in microstructure analysis (Davis and Smith, 1992; Spector et al., 1998), and the longer microstructure length, “***runs***”, as an exploratory analysis to determine the effects of microstructure length (Spector and St. John, 1998; Spector et al., 1998; Lin et al., 2013; Naneix et al., 2020). For each microstructure classification type, we determined the number of events, the duration of each event, and the number of licks per microstructure event. We also calculated the number and length of pauses after each microstructure event type, separated into 0.25-0.5s, 0.5-1s, 1-10s, 10-300s, and over 300s (Spector et al., 1998).

The estimated volume drank, calories drank, g/kg of alcohol, and lick sizes were calculated for each session using bottle weight differences, animal weight, and total number of validated licks, shown in **Fig. S1**. First, the average of all individual lick sizes (bottle difference divided by total licks) within 4-10 μL was used to calculate the daily lick size average for each daily cohort, to avoid inclusion of data from spilled bottles or any remaining artifactual lickometer counts. Then each rat’s estimated total volume based on licking was calculated using the daily average lick size for the cohort and that rat’s total licks. The estimated total volume or the recorded difference in bottle volume, whichever was lower, was divided by the total licks for the final individual rat’s lick size that day. Grand mean lick size was determined for each sex by the average of all final individual lick sizes, including all cocktails. This value was used to calculate all other volumes drank based on total number of licks per session.

Validated lickometer timestamps, preprocessed microstructure, estimated lick volumes, were migrated with another custom KNIME script to a relational database that also contained experimental variables including the rat’s identification, sex, and daily session information. Raw data from this database was queried using SQLite for further statistical analyses using GraphPad Prism 9/10.

Data from the last 4 sessions of each week was used for analyses. As the first day of drinking after a weekend often results in slightly different drinking responses (Samson, 2000), we by default did not analyze these days. In limited circumstances (1.3% of all sessions) for technical reasons such as spilled bottles or obstructed lickometer beams, we replaced individual sessions with the first day of the week, after the experimental parameters were reviewed and a consensus was reached by at least two experimenters.

Statistical analyses for overall drinking and microstructure events followed a standard strategy **(Fig. 1b)**.

First, an overall three-way Mixed Effects Analysis was used to determine if there was a sex difference in the compared fixed effects of sugar, sex, and alcohol percent, as well as interactions for each drinking and microstructure event. For clarity, all statistics from three-way analyses are found in **Table S1**, and the presence or absence of sex differences are noted in the text. If there were no sex differences, female and male groups were combined for further comparisons, otherwise the data were analyzed separately. In analyses of pause durations, sex differences were found for fructose and glucose cocktails separately. With sex differences identified within glucose, but not fructose cocktails, the data were analyzed across three groups: glucose males, glucose females, and fructose males and females (combined).

Next, two-way Mixed Effects Analyses were used to determine sugar and alcohol concentration differences, or pause duration and alcohol concentration differences. All two-way analyses with alcohol concentration as a factor resulted in a fixed effect of alcohol concentration, so for clarity, all statistics for alcohol concentration fixed effects are found in **Table S2**. For mixed-models analyses, we used a compound symmetry covariance matrix, fitted using restricted maximum likelihood (REML). *Post-hoc* comparisons used the Šídák method for glucose versus fructose comparisons and for comparing male and female glucose groups to a combined fructose group.

As there is a temporal delay between ingestion and entrance of glucose into the brain, the first 30 minutes of drinking behavior is often thought to be predominantly regulated by taste (Davis, 1973; Ackroff and Sclafani, 1991). For all microstructure analyses, all cocktail pairs with significant *post-hoc* comparisons were analyzed in 30-minute bins, to compare the first 30 minutes with the remaining seven 30-minute bins in the rest of the session to evaluate how measures of palatability or post-ingestive effects change during the session in conjunction with the other independent variables.

Statistical analyses for pauses of different lengths between microstructure events were conducted in the same way, except different lengths of pauses and sex differences were compared within each cocktail by three-way Mixed Effects Analyses, then pause durations and sugars were compared within each alcohol percent with two-way repeated measures (RM) Mixed Effects Analyses. The number of pauses is equivalent to the number of each microstructure, but the value for pauses is 1 less than the value of the number of microstructures total. All statistics reported for total pause number are identical to the statistics for microstructure number. Run pauses begin at 1-10s as the microstructure is longer than for cluster pauses.

## Results

### Total volume of cocktail drank during the session varies by sugar, alcohol concentration, and sex

We found that the volume of cocktail drank in the session varied depending on the sugar type, alcohol concentration, and sex. For clarity, all overall 3-way analyses and 2-way alcohol statistics are detailed in **Table S1** and **Table S2**. As there was a sex interaction in the overall 3-way analysis (**Table S1**), we analyzed female and male rats separately and compared the sexes within each sugar type. In both female and male rats, a two-way RM Mixed Effects Analysis revealed a fixed effect of *sugar type* (females: **Fig. 2a**, *F*_1,30_=5.107, *p*=0.0313; males: **Fig. 2b**, *F*_1,30_=14.74, *p*=0.0006), and an *alcohol concentration* x *sugar type* interaction (females: *F*_3,90_=11.55, *p*<0.0001; males: *F*_3,90_=43.57, *p*<0.0001). For both sexes, the volume of cocktail drank decreased generally as alcohol concentration increased. Overall, both male and female rats drank significantly more volumes of the glucose than fructose cocktails at the lower percentages of alcohol (1.25% and 2.5%). At 5% alcohol, both sexes drank either type of cocktail equally. At the highest concentration of alcohol (10%), only the male rats drank significantly more of the fructose than glucose containing cocktail.

**Figure 2.**
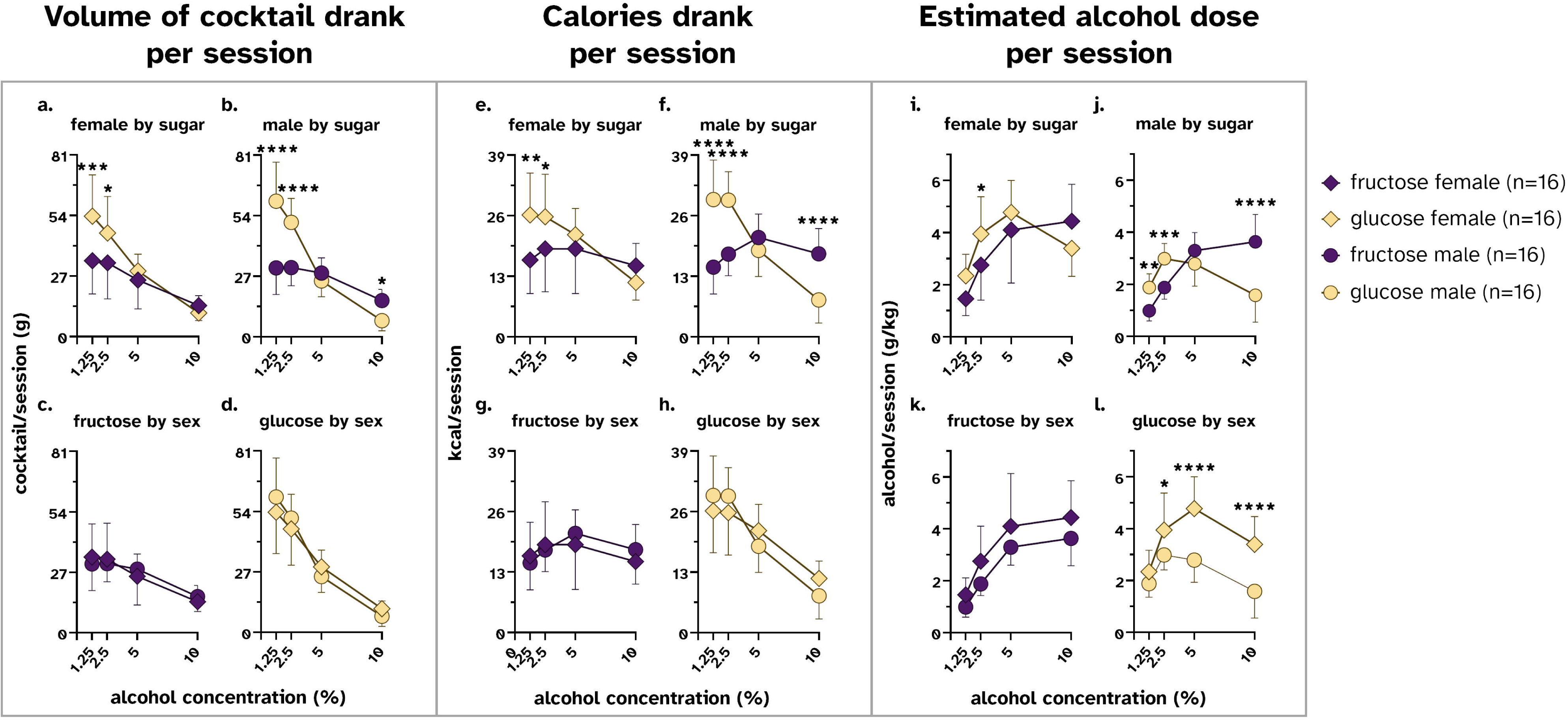
Estimated total volume, calories, and total alcohol dose (g/kg) drank vary by cocktail type, alcohol concentration, and sex. All data are shown as mean ± SD. **(a, b)** The volume drank of glucose cocktails was higher than fructose cocktails at the two lowest concentrations for both sexes. Male rats drank more fructose cocktail at the highest alcohol concentration. **(c, d)** Comparing the same data by sex show no significant differences within each sugar cocktail type, although there is an overall interaction in glucose cocktails. **(e, f)** Both females and males drank more calories per session of the glucose cocktail at the two lowest alcohol percentages, only male rats consumed more calories in the fructose cocktail at the highest alcohol concentration. **(g, h)** There were no significant sex differences in calories consumed within each cocktail type, although there is an overall interaction in glucose cocktails so that males showed a greater change in calories consumed across alcohol concentrations. (**i, j**) The estimated alcohol dose (g/kg) was greater for glucose cocktails in female rats only at 2.5%. In male rats, the estimated alcohol dose was greater for glucose cocktails at the lowest two alcohol concentrations; at the highest alcohol percentage, the estimated alcohol dose was greater for the fructose cocktail. (**k, l**) No sex differences in alcohol dose per session were found in rats drinking fructose cocktails; female rats drinking glucose cocktails achieved a higher estimated alcohol dose than males in a concentration-dependent manner. *p < 0.05, **p<0.01, ***p<0.001, ****p<0.0001 across cocktail type at a given alcohol concentration by Šídák’s multiple comparison test.

Within each sugar type, only the glucose cocktail showed sex differences in the volume consumed. Neither sugar type had a fixed effect of *sex*, and while there was no interaction when rats drank fructose (**Fig. 2c**), glucose cocktails showed an *alcohol concentration* x *sex* interaction. This meant that overall, at low alcohol concentrations, males tended to drink more glucose cocktails than their female counterparts, with the opposite trend at the highest alcohol concentration (**Fig. 2d**, *F*_3,90_=2.894, *p*=0.0396).

### Total calories drank during the session varies by sugar, alcohol concentration, and sex

As with volume, we found that the calories (kcal) consumed by the rats varied depending on the sugar type, alcohol concentration, and sex. As there was a sex interaction in the overall 3-way analysis (**Table S1**), we analyzed female and male rats separately. A two-way Mixed Effects Analysis revealed no fixed effect of *sugar type* in female rats (**Fig. 2e**), but this fixed effect was found in males (**Fig. 2f**, *F*_1,30_=5.092, *p*=0.0315), and there was an *alcohol concentration* x *sugar type* interaction in both sexes (females: **Fig. 2e**, *F*_3,90_=11.60, *p*<0.0001; males: **Fig. 2f**, *F*_3,90_=48.79, *p*<0.0001). At the two lowest concentrations of alcohol (1.25%, 2.5%), both male and female rats consumed more calories when drinking the glucose cocktail than the fructose cocktail. At the highest percentage of alcohol (10%), only male rats consumed more calories by drinking the fructose cocktail than the glucose cocktail.

Comparing calories drank between the sexes within each sugar type revealed modest sex differences depending on alcohol concentration. While there was no fixed effect of sex, each sugar type displayed an *alcohol concentration* x *sex* interaction; male rats showed a greater change in calories consumed due to alcohol concentration comparatively than female rats (fructose: **Fig. 2g**, *F*_3,90_=2.754, *p*=0.0476; glucose: **Fig. 2h**, *F*_3,90_=4.284, *p*=0.0071).

### Total estimated dose of alcohol drank during the session varies by sugar, alcohol concentration, and sex

We then estimated the alcohol dose consumed by rats during the session by accounting for their body weights during each session. Due to the 3-way sex interaction (**Table S1**), we analyzed female and male rats separately. There was a concentration effect of alcohol on the amount of alcohol consumed by rats; this differed by sugar type and sex. A two-way RM Mixed Effects Analysis revealed an *alcohol concentration* x *sugar type* interaction so that the alcohol dose achieved by the rats depended on the sugar type (females: **Fig 2i**, *F*_3,90_=12.10, *p*<0.0001; males: **Fig 2j**, *F*_3,90_=45.72, *p*<0.0001). Multiple comparison tests revealed that sugar type influenced g/kg alcohol intake only at 2.5% alcohol in female rats. However, male rats achieved a higher dose of alcohol when drinking the glucose cocktail than the fructose cocktail when the alcohol concentration was low (1.25%, 2.5%). This relationship was reversed when the alcohol concentration was high (10%).

When evaluating sex differences in g/kg alcohol intake within each sugar type, we found significant differences across sex in glucose but not fructose cocktails. A two-way Mixed Effects Analysis revealed a fixed effect of *sex* (fructose: **Fig 2k**, *F*_1,30_ =4.64, *p*=0.0394; glucose: **Fig 2l**, *F*_1,30_ =24.97, *p*<0.0001), as well as an *alcohol concentration* x *sex* interaction in rats drinking glucose cocktails only (*F*_3,90_=7.297, *p*=0.0002). Within fructose cocktails, the alcohol dose increased as the alcohol concentration in the cocktail increased, whereas in glucose cocktails the alcohol dose peaked at 5% and dropped off for both sexes, resulting in an inverted “U” shaped dose response pattern. Seen in both sugar types, regardless of alcohol concentration, was an elevated dose response in female rats compared to males, although this effect was greater with glucose cocktails. Multiple comparison tests within rats drinking glucose cocktails revealed that female rats achieved a significantly greater alcohol dose between 2.5% and 10% alcohol.

### Licks per cluster or run varies by the microstructure length and time interval within the session

To investigate the influence of sex, sugar type, and alcohol concentration on changes in palatability indices, we analyzed the number of licks within two microstructure categories: *clusters* (ILI >500ms) and *runs* (ILI >1s). Glucose achieves peak levels in the brain around 20-30 minutes after being consumed (Wakabayashi and Kiyatkin, 2015), therefore effects on licking microstructure that occur before this time during the session may be mediated less by rising glucose levels, while later changes in drinking microstructure may be influenced more strongly by direct effects of the sugar within the cocktail on the brain. To assess this, we analyzed each microstructure category separately for a) the first 30 minutes and b) from 30 minutes to 4 hours (i.e., the rest of the session).

There was an effect of *cocktail type*, and *alcohol concentration*, but not *sex* with ***licks per cluster*** (see **Table S1** for all 3-way analyses and **Table S2** for all 2-way fixed effects of alcohol). Due to the lack of a sex difference, females and males were combined for further analysis. For the first 30 minutes of the session, both fructose and glucose cocktails had a decrease in licks per cluster as alcohol concentration increased. A two-way RM Mixed Effects Analysis revealed an *alcohol concentration* x *sugar type* interaction, (**Fig. 3a**, *F*_3,186_=11.04, *p*<0.0001). Multiple comparison tests revealed significantly more licks per cluster only at the lowest alcohol concentration of 1.25% when rats were drinking the glucose cocktail. During the rest of the session, the same pattern remained: a two-way Mixed Effects Analysis revealed an *alcohol concentration* x *sugar type* interaction (**Fig. 3b**, *F*_3,186_=11.43, *p*<0.0001). Follow up *post-hoc* multiple comparison testing revealed that rats had greater licks per cluster when drinking the glucose cocktail with 1.25% alcohol than the equivalent fructose cocktail.

**Figure 3.**
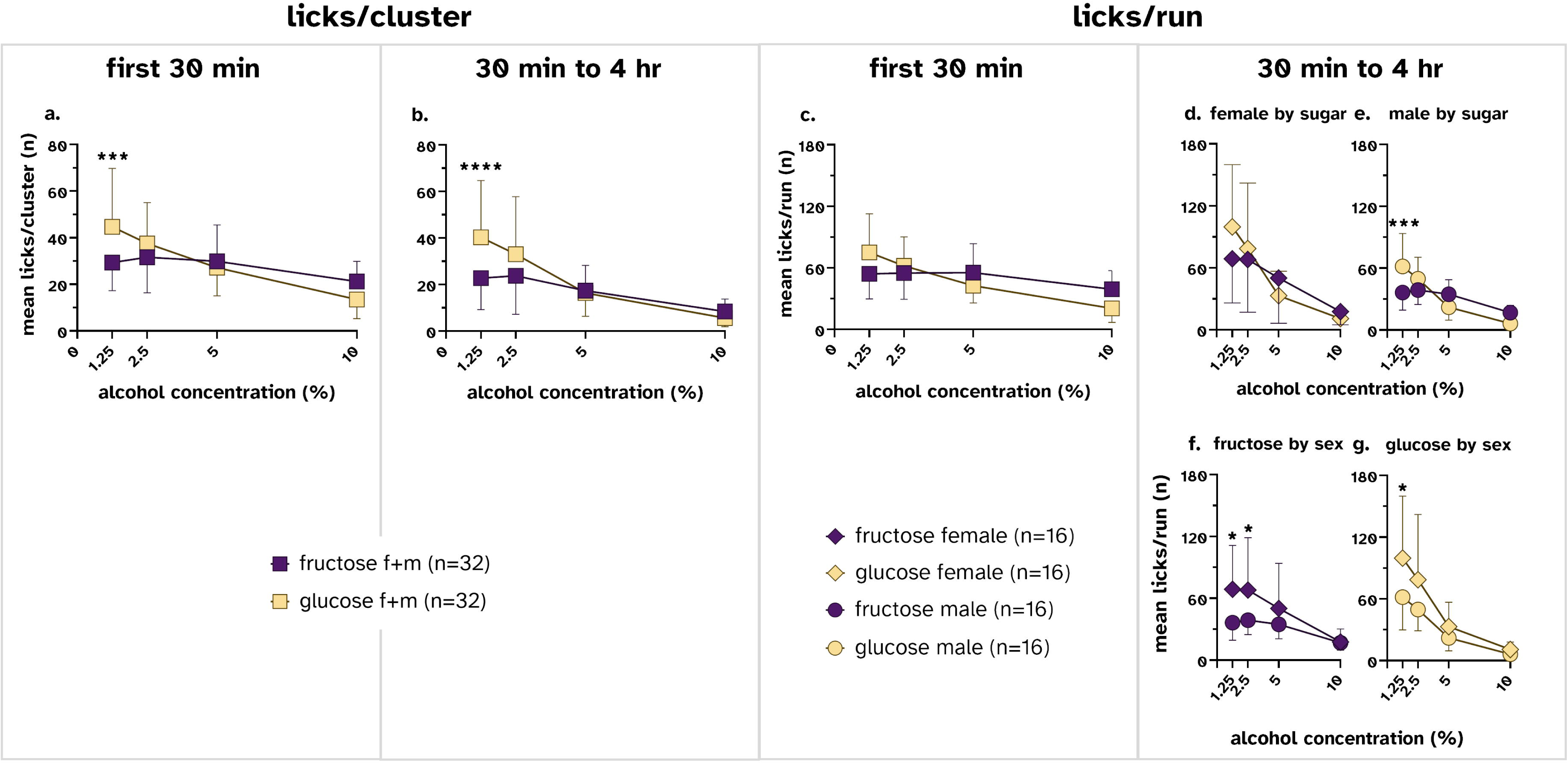
Licks per cluster or run depends on cocktail type, alcohol concentration, sex, and session time interval. All data are shown as mean ± SD. **(a)** Licks per cluster (ILI > 500 ms) for the first 30 minutes decreased for both cocktail types as alcohol concentration increased, with licks per cluster significantly greater with glucose cocktails than fructose at the lowest alcohol concentration. The same pattern of drinking was seen for **(b)** the rest of the session. **(c)** Licks per run (ILI > 1s) during the first 30 min of the session showed a similar overall interaction between cocktail type and alcohol concentration to licks per cluster, although no post-hoc differences between cocktail type were found. **(d, e)** For the rest of the session, while both sexes reduced the number of licks per run as alcohol concentration increased, sex differences were found. **(e)** Only male rats had a significantly higher number of licks per run drinking the lowest alcohol concentration glucose cocktail compared to the fructose cocktail. (**f**) During this part of the session, female rats drinking fructose cocktails had greater licks per run than males for the two lower percentages of alcohol. (**g**) Female rats also had greater licks per run compared to male rats during the later portion of the session when drinking the lowest alcohol concentration glucose cocktail. *p < 0.05, **p<0.01, ***p<0.001, ****p<0.0001 across sugar type or sex at a given alcohol concentration by Šídák’s multiple comparison test.

Evaluating the longer microstructure category ***licks per run***, there was a similar overall decrease in licks per run as alcohol concentration increased, which varied by sugar type and sex. During the first 30 minutes of the session, there was no fixed effect of *sex* or any sex interactions. Combining the sexes, a two-way RM Mixed Effects Analysis revealed a *sugar type* x *alcohol concentration* interaction, (**Fig. 3c**, *F*_3,186_=14.06, *p*<0.0001), such that the licks per run for glucose cocktails were higher than the fructose cocktail at lower concentrations of alcohol. At higher concentrations, this relationship was reversed, but *post-hoc* multiple comparisons found no differences at any specific alcohol concentration.

During the rest of the 4-hr session, there were sex differences in the licks per run, so each sex was analyzed separately. There was an *alcohol concentration* x *sugar type* interaction for both sexes (female: **Fig. 3d**, *F*_3,90_=3.281, *p*=0.0245; male: **Fig. 3e**, *F*_3,90_=11.80, *p*<0.0001).

Only male rats drinking the glucose cocktail had more licks per run at the lowest alcohol percentage than male rats drinking the equivalent fructose cocktail. This difference was not observed in female rats. Analyses within cocktail type revealed fixed effects of *sex* (fructose: **Fig. 3f**, *F*_3,90_=5.159, *p*=0.0305; glucose: **Fig. 3g**, *F*_3,90_=5.892, *p*=0.0214) on licks per run.

Further, only in rats drinking the fructose cocktail was there an *alcohol concentration* x *sex* interaction (*F*_3,90_=3.974, *p*=0.0104). Female rats had a significantly greater number of licks per run than male rats when drinking fructose cocktails containing the two lowest alcohol concentrations. Female rats also had greater licks per run during this session interval when drinking the lowest alcohol concentration glucose cocktail.

### The number of pauses after a cluster or run varies between early and late intervals within the session

#### Cluster pause number has stronger sugar differences for male rats in the first 30 minutes

We first compared all groups within each time interval for the number of pauses after a cluster (ILI > 500 ms, “*cluster pauses*”) to determine sex differences (statistics detailed in **Table S1** for all 3-way analyses and **Table S2** for all 2-way fixed effects of *alcohol concentration*).

#### Cluster pauses vary strongly by sex and sugar in the first 30 minutes

During this interval, we found sex differences within the overall 3-way analysis, and therefore kept the sexes separate in further analyses. While female rats showed no effect of sugar (**Fig. 4a)**, male rats showed a significant *sugar type* x *alcohol concentration* interaction (two-way RM Mixed Effects Analysis, **Fig. 4b**, *F*_3,90_=10.61, *p*<0.0001), indicating that the number of post-cluster pauses depended on the type of cocktail as the alcohol concentration increased. Indeed, *post-hoc* multiple comparison tests revealed that male rats had more post-cluster pauses when drinking 10% alcohol fructose cocktails compared to the same alcohol concentration glucose cocktail. When looking at within-sugar comparisons by sex, we saw a greater number of differences between male and female rats drinking the glucose cocktail. Rats drinking the glucose cocktail had a significant fixed effect of *sex* (**Fig. 4d**, *F*_1,90_=4.731, *p*=0.0376), and a significant *alcohol concentration* x *sex* interaction (*F*_3,90_=5.776, *p*=0.0012). Whereas most differences seen thus far have been restricted to either the lowest or highest percentage of alcohol, *post-hoc* multiple comparisons revealed a significant difference between male and female rats drinking the glucose cocktail at the 5% alcohol percentage. No significant fixed effects nor interactions were seen in rats drinking fructose (**Fig. 4c**).

**Figure 4.**
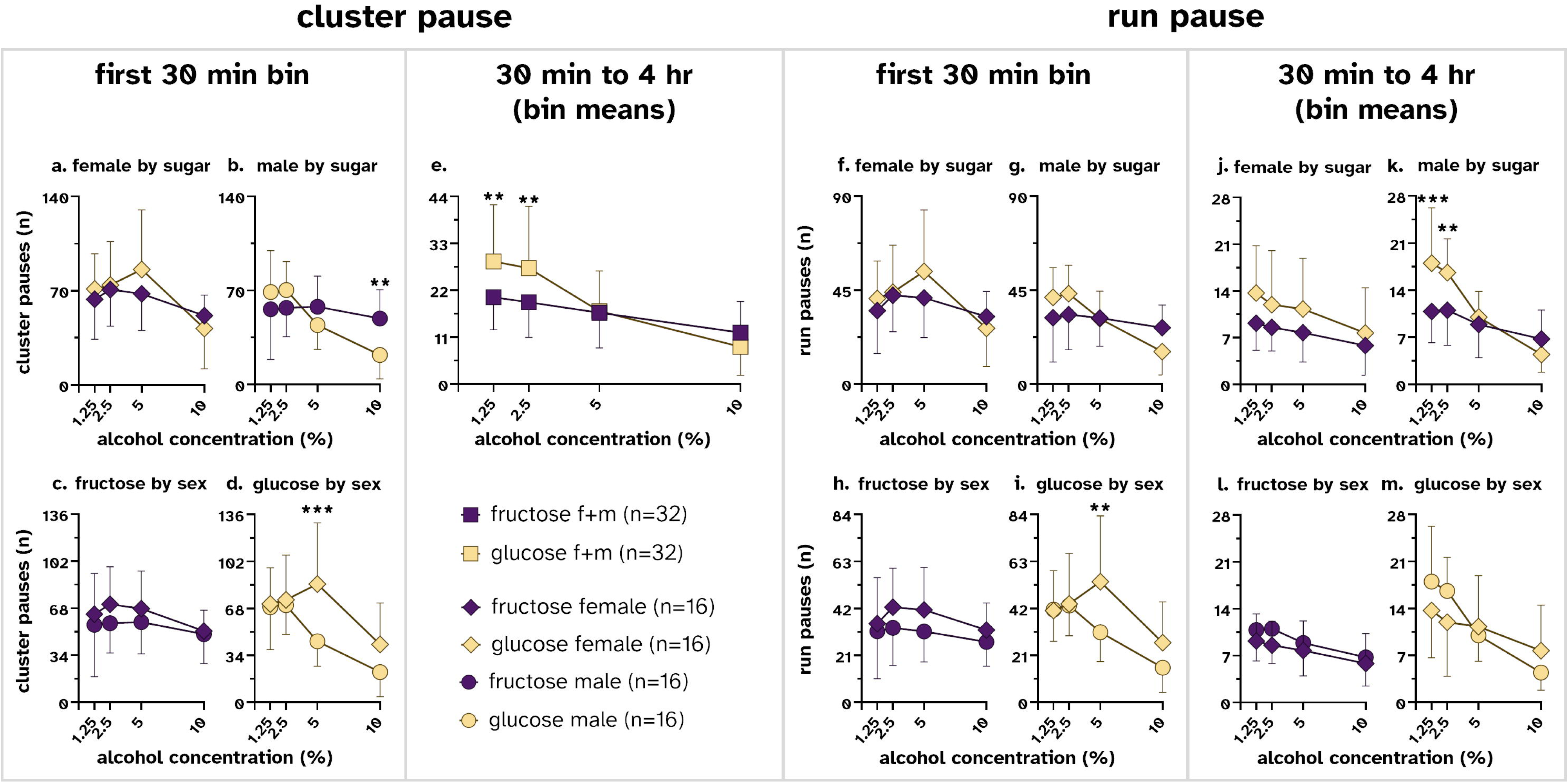
The number of pauses varies by microstructure type and when during the session they occur. All data are shown as mean ± SD. **(a, b)** The number of pauses after a cluster (ILI > 500 ms) for the first 30 minutes of the session were significantly higher for male rats when drinking the highest alcohol concentration fructose cocktail compared to the glucose cocktail. **(c)** When evaluated by sugar type, there were no sex differences in fructose cocktails. **(d)** There was a sex difference in glucose cocktails where female rats had significantly more post-cluster pauses than male rats at 5% alcohol. **(e)** There were no sex differences for the remainder of the session; rats overall had significantly more pauses after a cluster while drinking 1.25% and 2.5% alcohol glucose cocktails. (**f, g**) The number of pauses after a run (ILI > 1s) for the first 30 min of the session showed no mean differences between cocktail types for either sex. (**h, i**) Within each cocktail type, female rats had a higher number of post-run pauses compared to male rats in glucose, but not fructose, 5% alcohol cocktails. (**j, k**) For the remainder of the session, only male rats showed a greater number of pauses after a run when drinking glucose cocktails at the two lowest concentrations of alcohol; there were no differences between cocktails for female rats. (**l, m**) When comparing by cocktail type, neither sex had significantly different post-run pauses when drinking either cocktail type. *p<0.05, **p<0.01, ***p<0.001, ****p<0.0001 across cocktail type or sex at a given alcohol concentration by Šídák’s multiple comparison test.

#### The number of cluster pauses show no sex differences for the remainder of the session

There were no sex differences in the number of post-cluster pauses for the remainder of the session, so the data from both sexes were combined. Post-cluster pause number decreased over a wider range across alcohol concentration for the glucose cocktail (*sugar type* x *alcohol concentration* interaction, **Fig. 4e***, F*_3,186_=11.87, *p*<0.0001). Specifically, there were more post-cluster pauses at the two lowest concentrations of alcohol when rats were drinking the glucose cocktail compared to the fructose cocktail.

### Run pause number shows stronger sugar differences for male rats after 30 minutes

Next, we investigated the number of pauses after a run (ILI > 1s; “*run pauses*”) for each group. Each group was compared within each time interval to determine sex differences (statistics detailed in **Table S1** for all 3-way analyses and **Table S2** for all 2-way fixed effects of *alcohol concentration*).

#### Run pauses show significant effects of sex but not sugar in the first 30 minutes

Sex differences were detected during the first 30 minutes of the session and therefore we kept the sexes separate for the next analyses. When examining the number of post-run pauses when drinking different cocktail types and alcohol concentration, we saw a similar alcohol concentration-dependent pattern as seen with post-cluster pauses. A two-way RM Mixed Effects Analysis revealed no fixed effect of *sugar type* in either sex. There was no overall significant *alcohol concentration* x *sugar type* interaction in females (**Fig. 4f**), though it was present in male rats (**Fig. 4g**; *F*_3,90_=8.443, *p*<0.0001). *Post-hoc* multiple comparisons revealed no individual differences between run pauses between cocktail types at any alcohol percentage for male rats.

When comparing the number of run pauses within each cocktail type between male and female rats, we found no fixed effect of *sex* in either sugar type. There was a significant *alcohol type* x *sex* interaction in rats consuming glucose cocktails (**Fig. 4i**, *F*_3,90_=4.486, *p*=0.0056), but not in fructose cocktails (**Fig. 4h**). *Post-hoc* multiple comparisons revealed that the 5% alcohol concentration was significantly different between female and male rats.

#### Run pauses show sex differences in the remainder of the session

Due to the presence of sex differences in the omnibus test, the post-run pauses of male and female rats were analyzed separately for this time portion of the session. Male rats (**Fig. 4k**) showed an *alcohol concentration* x *sugar type* interaction (*F*_3,90_=11.79, *p*<0.0001) while female rats (**Fig. 4j**) did not have any effect of *sugar* or interaction. *Post-hoc* multiple comparisons revealed that male rats had a significantly greater number of post-run pauses when drinking glucose cocktails at the two lowest concentrations of alcohol.

When examining the number of post-run pauses within each cocktail type between male and female rats for this part of the session, we saw no fixed effect of *sex* in either sugar type.

However, there was a significant *alcohol concentration* x *sex* interaction in rats consuming the glucose (**Fig. 4m**; *F*_3,90_=6.024, *p*=0.0009) but not the fructose cocktail (**Fig. 4l**). Despite an overall interaction, *post-hoc* multiple comparisons did not detect any concentration-specific differences when rats were drinking the glucose cocktail.

### Cocktail differences in the number of short post-microstructure pauses depend on when they occurred in the session

To further assess the influence of session time on pauses in drinking, we examined the number of post-cluster and -run pauses of different durations between the two cocktails during the first 30 minutes of the session compared to the rest of the session across each alcohol concentration. For each type of drinking microstructure (i.e. a *cluster* or *run*), we examined how long the pauses were by quantifying how many fell into five different bins of increasing duration (0.5s-1s, 1-10s, 10-300s, and longer than 300s), for the whole session.

In order to analyze different lengths of pauses between the first 30 minutes and the remainder of the session, we first identified which cocktail types displayed sex differences (all 3-way analysis results are found in **Table S1**). For both post-cluster (ILI > 500 ms) and post-run (ILI > 1s) pauses, there were sex differences within the glucose cocktails, so glucose groups were kept separate between the sexes, and there were not sex differences within the fructose groups, so data from the two sexes were combined into one fructose group.

We then analyzed each sex and cocktail type in a two-way analysis of the entire session and found that every group varied strongly across both alcohol concentration and pause duration (**Table S3**), so we then examined effects of cocktail type and pause duration within each alcohol concentration separately (**Fig. S3, Table S4**). We found significant cocktail type differences at pause durations of 0.5-1s and 1-10s for post-cluster pauses and pause durations of 1-10s only for post-run pauses. Therefore, for the following analyses, we only analyzed these shorter pause durations across the two time periods within the session.

#### At lower alcohol concentrations, the number of short post-cluster pauses only differs between male glucose cocktail drinkers and fructose cocktail drinking rats of both sexes after 30 minutes

First, we analyzed the number of post-cluster (ILI > 500 ms) pauses across early (first 30 min) and remaining (30 min to 4 hr) time periods during the session. As the number of post-cluster pauses that were 0.5-1s long at the two lowest alcohol concentrations were not significantly different over the entire session (**Fig. S3a-d**), we only analyzed 1-10s post-cluster pauses. For the first 30 minutes of the session, rats drinking 1.25% and 2.5% alcohol showed no significant differences in 1-10s long post-cluster pauses between rats drinking glucose or fructose cocktails (**Fig. 5a, c**). For the remainder of the session, significant main effects were found in rats drinking 1.25% (**Fig. 5b**, one-way ANOVA, *F*=9.381, *p*=0.0003) and 2.5% alcohol (**Fig. 5d**, *F* = 7.756, *p* = 0.001). Subsequent *post-hoc* multiple comparisons revealed that male rats drinking glucose cocktails paused for 1-10s more often after a cluster than rats drinking fructose cocktails. At both 1.25% and 2.5% alcohol concentrations, pauses for female rats drinking the glucose cocktail were not different than for the fructose-drinking rats.

**Figure 5.**
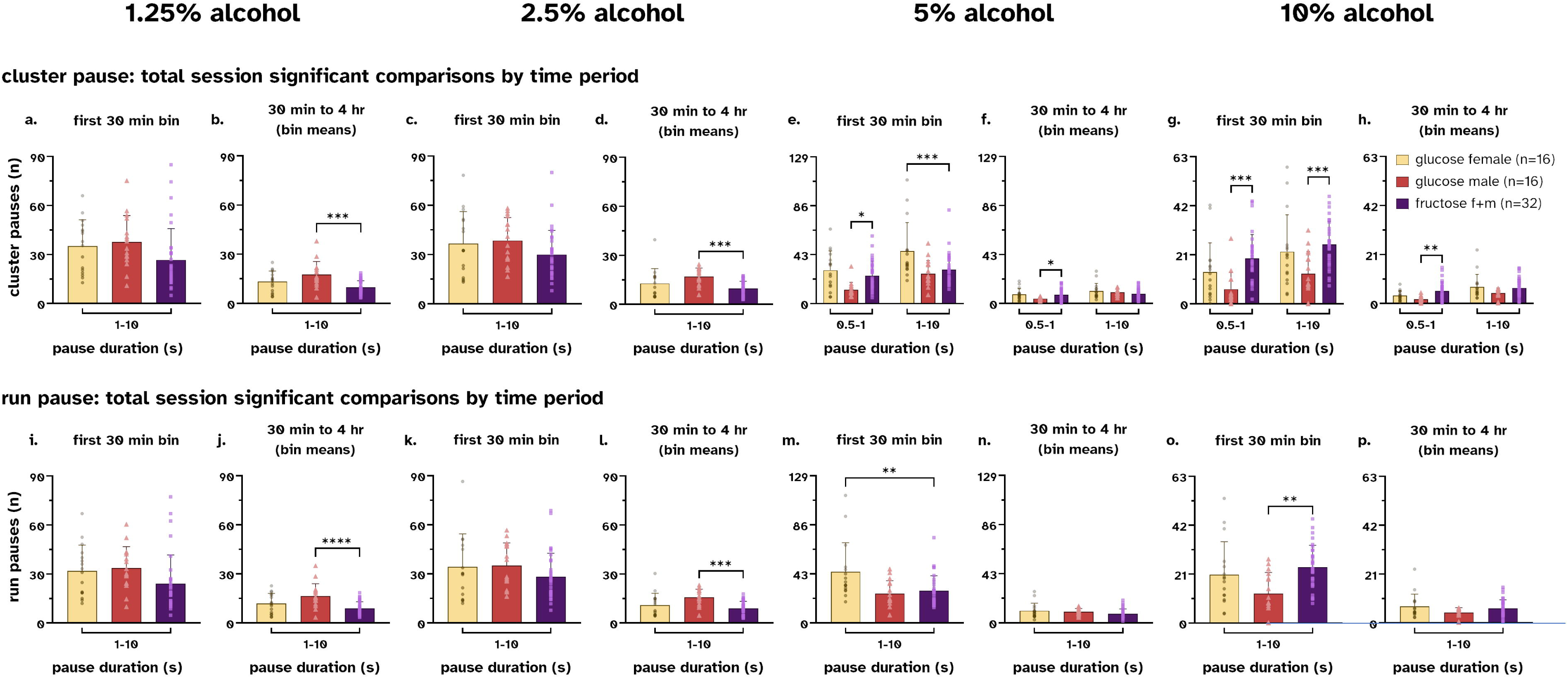
Sugar differences in cluster and run pause numbers depend on time interval within the session. **(a)** During the first 30 minutes of the session, the number of post-cluster pauses that were 1-10s at 1.25% alcohol was similar between rats drinking either the fructose or glucose cocktails. **(b)** For the remainder of the session, on average male rats drinking the glucose cocktail paused after a cluster for 1-10s significantly more often than rats drinking the fructose cocktail. **(c)** During the first 30 minutes of the session, the number of post-cluster pauses at 2.5% alcohol were not different across cocktail drinking groups. **(d)** For the remainder of the session, similar to 1.25% alcohol, male rats drinking the 2.5% alcohol glucose cocktail had significantly more post-cluster pauses than rats drinking the fructose cocktail. **(e)** During the first 30 minutes, male rats drinking the 5% alcohol glucose cocktail showed significantly fewer 0.5-1s pauses compared to rats drinking the fructose cocktail. Female rats showed a greater number of 1-10s pauses compared to rats drinking the fructose cocktail. **(f)** For the rest of the session, 0.5-1s post-cluster pauses at 5% alcohol were lower for glucose cocktail-drinking male rats compared to rats drinking the fructose cocktail. **(g)** At 10% alcohol, male rats drinking the glucose cocktail had significantly fewer 0.5-1s and 1-10s long pauses than rats drinking the fructose cocktail. **(h)** During the remainder of the session, only male rats showed significantly fewer 0.5-1s post-cluster pauses compared to fructose cocktail drinking rats. **(i, k)** During the first 30 minutes of the session, rats drinking cocktails at 1.25% and 2.5% had no significant differences in the number of post-run pauses that were 1-10s long. **(j, k)** During the remainder of the session, male rats drinking glucose cocktails with 1.25% or 2.5% alcohol on average paused for 1-10s significantly more often after a run than rats drinking the equivalent alcohol fructose cocktail. **(m)** At 5% alcohol, during the first 30 minutes of the session, female rats drinking the glucose cocktail paused for 1-10s significantly more than rats drinking the equivalent fructose cocktail. **(n)** For the remainder of the session, all groups had low run pauses 1-10s long. **(o)** At 10% alcohol, rats drinking the fructose cocktail paused more often compared to male rats drinking the glucose cocktail in the first 30 minutes of the session. **(p)** For the remainder of the session, both sugar types had low run pauses 1-10s long. *p<0.05, **p<0.01, ***p<0.001, ****p<0.0001 comparing the female glucose or male glucose group to the female+male fructose group at a given pause duration by Šídák’s multiple comparison test.

At the 5% and 10% alcohol concentrations, the number of post-cluster pauses were significantly different over the entire session when they were either 0.5-1s and 1-10s long (**Fig. S3c, d**), thus both pause durations were subsequently analyzed. At 5% alcohol, in the first 30 minutes of the session, there was a significant fixed effect of *pause duration* (**Fig. 5e**, two-way RM Mixed Effects Analysis, *F*_1,61_=51.30, *p*<0.0001), a fixed effect of *sugar type* (*F*_2,61_=7.521, *p*=0.0012), and ultimately a *pause duration* x *sugar type* interaction, *F*_2,61_=5.281, *p*=0.0077).

*Post-hoc* multiple comparison analysis revealed that male rats drinking glucose cocktails paused for 0.5-1s significantly less often than their fructose cocktail drinking counterparts. On the other hand, female rats paused significantly more often for 1-10s than fructose cocktail drinking rats. For the remainder of the session at the 5% alcohol percentage, there was a fixed effect of *pause duration* (**Fig. 5f**, *F*_1,61_=30.54, *p*<0.0001) and a *pause duration* x *sugar type* interaction (*F*_2,61_=7.281, *p*=0.0015). *Post-hoc* multiple comparisons revealed that male rats drinking glucose cocktails continued to pause for 0.5-1s less often than their fructose cocktail drinking equivalents, and that for longer 1-10s, no significant group differences were found at this session interval. At 10% alcohol, in the first 30 minutes of the session, there was a fixed effect of *pause duration* (**Fig. 5g**, *F*_1,61_=31.16, *p*<0.0001), a fixed effect of *sugar type* (*F*_2,61_=9.172, *p*=0.0003), and no significant interaction. *Post-hoc* multiple comparisons highlighted that male rats drinking the glucose cocktail paused for 0.5-1s and 1-10s less often than rats drinking the fructose cocktail. In the later portion of the session, there was a fixed effect of *pause duration* (**Fig. 5h**, *F*_1,61_=35.38, *p*<0.0001), as well as an overall fixed effect of *sugar type* (*F*_2,61_=4.520, *p*=0.0148), and a significant *pause duration* x *sugar type* interaction (*F*_2,61_=3.251, *p*=0.0456). Similar to the first 30 minutes of the session, *post-hoc* multiple comparisons revealed that male rats drinking the glucose cocktail in the later portion of the session paused for 0.5-1s less often than rats drinking the fructose cocktail. Unlike the first 30 minutes, no significant within-group differences were observed at longer pause intervals (1-10s) during this part of the session and alcohol concentration.

#### Run pauses show sugar differences after 30 minutes only for male rats at low alcohol concentrations and opposite sugar differences for both sexes at higher alcohol concentrations

Next, we analyzed the number of pauses after a run (ILI > 1s, “run pauses”) during the first 30 minutes of the session and then during the rest of the session as with pauses after a cluster. Unlike clusters, only pause lengths of 1-10s after a run had significant sugar differences (**Fig. S3e-h**), therefore only this pause duration was analyzed.

At both the 1.25% and 2.5% alcohol concentration, there were no differences in the number of times a rat paused after a run when drinking either sugar cocktail during the first 30 minutes of the session (**Fig. 5i, k**). However, for the remainder of the session, there were significant main effects for the 1.25% (**Fig. 5j**, one-way ANOVA, *F_2,61_*=9.766, *p*=0.0002) and 2.5% alcohol concentration (**Fig 5l**, *F*_2,61_=8.661, *p*=0.0005). *Post-hoc* multiple comparisons revealed that at both alcohol concentrations, male rats drinking the glucose cocktail paused for 1-10s more often than the fructose-drinking rats.

During the first 30 minutes of the session at 5% alcohol (**Fig. 5m**), there was a main effect (one-way ANOVA, *F_2,61_*=6.607, *p*=0.0025). Specifically, female rats drinking the glucose cocktail paused after a run significantly more than the fructose cocktail drinking rats. In the later portion of the 5% alcohol session (**Fig. 5n)**, there was no effect of *sugar type*.

During the first 30 minutes of the 10% alcohol session (**Fig. 5o**), there was a main effect (one-way ANOVA, *F_2,61_*=6.114, *p*=0.0038). Here, male rats drinking the glucose cocktail paused less often for 1-10s after a run than the fructose cocktail drinking rats. During the rest of the session (**Fig. 5p**), there was again no effect of *sugar type*.

## Discussion

The objective of this study was to determine how the addition of glucose or fructose to different concentrations of alcohol would impact drinking patterns in Wistar rats. We sought to test the hypothesis that glucose containing cocktails would have greater centrally mediated behavioral effects than the equally caloric but less brain-penetrant fructose. As part of our study objectives, we also determined if there were any sex differences in these drinking patterns. We found that rats of both sexes drank more lower alcohol concentration glucose cocktails than cocktails containing fructose by volume and by overall calories. As alcohol concentration increased, these differences in drinking either disappeared or in some cases reversed. When considering the dose of alcohol, adding glucose seemed to increase the sensitivity to the effects of alcohol by shifting the dose response curve leftward compared to fructose cocktails. We found that drinking microstructures traditionally associated with palatability remained stable for both types of cocktails over the entire drinking session. In contrast, indices of post-ingestive effects appeared to be more varied, showing greater influences of sex and the time within the session. Notably, rats of both sexes showed higher post-ingestive measures with glucose cocktails at lower concentrations of alcohol than rats drinking fructose cocktails, particularly during the time periods when post-drinking central regulation is assumed to be strongest.

However, as the alcohol concentration increased, the differences in some post-ingestive measures between the two types of cocktails either disappeared or in some cases reversed, although this was measure- and sex- dependent.

### Sugar type and alcohol concentration interact to impact cocktail intake

It has been long known that adding sweeteners like sucrose or saccharin will increase alcohol intake in rats over a range of different experimental parameters (Eriksson, 1969; Gilbert, 1974; Samson et al., 1996; Czachowski et al., 1999). Even though the use of a sucrose- or saccharin-substitution procedure (i.e., “sweetened-fade”, (Samson, 1986; Rassnick et al., 1992) likely models human alcohol drinking patterns closely and is widely used to train rats to drink alcohol, using sweeteners in preclinical research is considered somewhat problematic. This is in part because sweeteners could obscure whether alcohol drinking is mediated by the psychoactive effects of alcohol alone or by interaction with the sweet taste of the cocktail.

However, this may be an inaccurate distinction. Some sugars, mainly glucose, can rapidly cross the blood-brain barrier (BBB) via facilitated, gradient-dependent diffusion by the glucose transporter-1 (GLUT-1; de Vries et al., 2003; Fellows et al., 2006; Fellows & Boutelle, 1993; Silver & Erecińska, 1994). This rise in glucose concentration occurs at behaviorally relevant timescales and can be detected in the brain soon after a rat begins to drink glucose, or is administered glucose via other routes of administration, such as intravenously (Wakabayashi and Kiyatkin, 2015; Kuebler et al., 2022). Moreover, γ-aminobutyric acid (GABA), orexin/hypocretin, neuropeptide Y (NPY), and melanin-concentrating hormone (MCH) neurons, are glucose sensitive (Amoroso et al., 1990; Muroya et al., 1999, 2001; Burdakov et al., 2005). These neurons change their activity based on the extracellular glucose concentration (Routh et al., 2014). This glucose sensitivity could be linked to the development of experience-dependent escalation of glucose drinking over fructose (Ackroff and Sclafani, 1991; Wakabayashi et al., 2016), an equally caloric, natural monosaccharide sugar that crosses into the brain much more slowly (Duelli and Kuschinsky, 2001). Therefore, alcoholic cocktails containing glucose monosaccharides may have distinct central, psychoactive effects from similarly sweet alcoholic cocktails containing fructose. This distinction is relevant to people because many commercially manufactured alcoholic beverages contain a mix of glucose and fructose monosaccharides in various proportions, ranging from 42% to 70% fructose versus glucose, with a 55% fructose, 45% glucose mixture often found in these beverages (Khorshidian et al., 2021).

During our experiment, we found that, by volume, both male and female rats drank more glucose-containing cocktails at the two lower concentrations of alcohol. As the alcohol concentration increased, we found no significant difference in the volume drank at the middle 5% concentration. Finally, at the highest concentration tested (10% alcohol), only male rats drank significantly more fructose- than glucose- containing cocktails (**Fig 2a-d**). Notably, no sex differences in volume drank within each cocktail type were seen. At a first look, it seems that both female and male rats prefer drinking glucose cocktails more than fructose cocktails when the alcohol concentration is low, and that this preference narrows, approaches parity, and then eventually reverses slightly as the alcohol percentage increases. This trend also continued when we considered the caloric value of the cocktails that the rats were consuming. Alcohol, glucose, and fructose are all nutritive and have caloric values (Atwater and Benedict, 1902; Ayoub et al., 2020; Khorshidian et al., 2021), and many glucose-sensitive neurons and circuits are implicated in energy homeostasis (Burdakov et al., 2005; Mergenthaler et al., 2013; Routh et al., 2014). However, it is clear from **Fig. 2e, f** that the amount of glucose or fructose compared to alcohol that the rats consumed in each cocktail was not directly related to the calories in that cocktail. For example, at 10% alcohol, both glucose and fructose cocktails have the same caloric value, yet male rats drank less of the glucose cocktail than the fructose cocktail. However, while rats of both sexes drank similar volumes of cocktails during these sessions (**Figs. 2c, d**) female rats were smaller than males throughout the experiment (**Fig S2**). Accounting for differences in body weight and considering the estimated dose of alcohol (g/kg) drank during the sessions, rats drinking glucose cocktails showed a significantly different alcohol dose response curve than rats that drank fructose cocktails. Namely, in rats of both sexes, there was a clear leftward shift in the alcohol dose response curve when glucose was added to the cocktails compared to fructose cocktails. Thus, one pharmacological interpretation of the behavior is that adding glucose made rats more sensitive to the effects of alcohol than the same concentration of fructose. As well, female rats drank more alcohol per body weight than male rats, supporting earlier findings that female rats drink more alcohol than male rats (Eriksson, 1969). One question to answer stemming from whole session behavior was: were these differences in drinking these cocktails due to differences in taste, or differences in their central effects?

### Palatability shows consistency or sex differences depending on the microstructure type

One frequently reported measure of taste is the number of licks per cluster, that is, licks that occur within a group of licks separated by an interlick interval (ILI) greater than 500 ms (Davis and Smith, 1992; Spector et al., 1998; Smith, 2001). Licks per cluster are positively correlated with palatability. For example, rats decrease their licks per cluster when drinking unpalatable solutions (Hsiao and Fan, 1993; Spector and St. John, 1998). Additionally, in male rats, “burst size”, an analogous behavioral measure to licks per cluster, decreases over the course of drinking sucrose solutions over a 1-hr session, suggesting diminishing palatability (Spector et al., 1998). In our current study, the number of licks per cluster for all cocktails and alcohol concentrations was largely consistent throughout the session **(Fig. 3a-b)**. Broadly, all glucose cocktail drinking rats had higher licks per cluster than rats drinking fructose cocktails when the alcohol concentration was low. As the concentration increased, this difference in the number of licks reduced until it reached parity at 5% alcohol and largely remained at 10% alcohol. Therefore, we interpret the changes in this behavioral measure as low alcohol concentration glucose cocktails are more palatable than fructose cocktails, and that this difference in palatability largely disappears once the alcohol content increases past 5%.

#### Importantly, we found that these drinking patterns were independent of the time within the session, suggesting that this index of the cocktail’s taste did not change for the rats as the session progressed

Others have also reported licks that occur within a group of licks separated by an ILI greater than 1s, (i.e. a “run”) as a complementary measure of palatability (Davis, 1989; Schier and Spector, 2016; Renteria et al., 2020). This allows for longer breaks within microstructures to be used to assess palatability. Rats drinking alcohol-free glucose solutions have greater licks per run when compared to those drinking the same concentration fructose solution; this has been interpreted as increased palatability of glucose over fructose solutions (Schier and Spector, 2016). As well, the number of licks per run have been reported to be sensitive to increases in alcohol concentration (Richter and Campbell, 1940; Eriksson, 1969; Samson, 1986), suggesting that this longer ILI measure is also able to detect changes in taste. When we analyzed our data using this longer interval, we found that there was a significant contribution of sex in the later portion of the session **(Fig. 3c-g)**. Namely, male rats changed the number of licks per run while drinking glucose cocktails more so than male rats drinking fructose cocktails in an alcohol concentration dependent manner (**Fig. 3e**). The differences in this measure of palatability between the cocktails was less robust in female rats (**Fig. 3d**). Moreover, female rats evinced greater palatability for both types of cocktails in an alcohol concentration-dependent manner compared to male rats in the later portion of the session (**Fig. 3f, g**). As well, female rats showed a greater range in this measure of palatability than male rats across the different alcohol concentrations **(Fig. 3d-g)**. One parsimonious interpretation of these data is that male rats more consistently discriminate between the palatability of the glucose and fructose cocktails at the lowest concentration of alcohol (**Fig 3e**). In contrast, female rats found glucose cocktails more palatable at lower concentrations of alcohol than male rats (**Fig. 3g**). Interestingly, these changes in taste appeared to be more pronounced in the later portion of the session, during a time when the cocktails have entered the brain and are influencing drinking behavior.

Taken together, these results suggest that this measure of palatability is detecting a centrally mediated component of taste, and further that this measure is sensitive enough to detect sex differences in this process. However, it should be noted that the construct validity of licks per run as a measure of palatability is less established than licks per cluster (Spector et al., 1998). Thus, careful future work will need to be undertaken to further determine the relationship between this behavioral measure, sex differences, and orosensory positive feedback when considering cocktail drinking.

### Post-ingestive effects vary across sex, time interval, and microstructure type

While the size of clusters (ILI > 500 ms) or runs (ILI > 1s) is tied to palatability and taste, the number of pauses between clusters or runs of licks is thought to reflect centrally mediated satiety (Davis, 1973; Davis and Smith, 1992; Spector et al., 1998; Naneix et al., 2020). This is in part because the number of pauses after a cluster or run of licks is related to how often the rat returns to drinking. As rats become increasingly satiated due to the greater central effects of a cocktail (e.g. the “post-ingestive load”), they initiate drinking bouts less often, leading ultimately to a decline in the number of pauses during a lengthy drinking session (Naneix et al., 2020). A critical component of this centrally mediated process is the time gap between the time a rat first begins to drink a cocktail, and the moment sufficient amounts of the cocktail arrive in the brain via the blood-brain barrier. Most microstructure drinking studies focus on the first thirty minutes of drinking where consumption is thought to be dynamically regulated initially by the sensory components of taste, then increasingly by post-ingestive feedback (Davis, 1989). Studies directly measuring both brain glucose and alcohol during voluntary drinking support this timeframe. For example, in male Long-Evans rats drinking a 10% glucose solution, increases in glucose are detected in the brain minutes after the start of drinking, reaching a peak approximately 20-30 minutes later (Wakabayashi and Kiyatkin, 2015). Moreover, alcohol levels in male Wistar rats drinking a 10% alcohol solution show similar pharmacokinetics, reaching a peak approximately 13-55 minutes after their first bout (Nurmi et al., 1999). Importantly, while both glucose and alcohol can enter the brain and have direct actions on central neurons (Nurmi et al., 1999; Duelli and Kuschinsky, 2001; Burdakov et al., 2005; Abrahao et al., 2017), fructose has no known direct central action on the brain over the same time frame.

In our current study, we measured the numbers of pauses between drinking bouts throughout the first 30 minutes and the remaining 3.5 hours of the session (**Fig. 4**). When considering the later portion of session where central effects of the cocktails would be the greatest, the average number of post-cluster pauses dropped across both cocktails for both sexes, presumably due to effects like satiety. Yet rats of both sexes drinking the glucose cocktail at the two lowest concentrations of alcohol clearly showed less satiety than rats drinking the fructose cocktail (**Fig 4e**). This difference in satiety between the two types of cocktails disappeared as the alcohol concentration increased. In contrast, during the first 30 minutes of the drinking session, when centrally mediated satiety effects are minimal, male rats showed significantly fewer cluster pauses when drinking the glucose cocktail than the fructose cocktail at the highest concentration of alcohol (**Fig. 4b**). Female rats, in comparison, did not show any significant differences in the number of post-cluster pauses between the glucose and fructose cocktails (**Fig. 4a**). When directly comparing glucose cocktail post-cluster pauses at the beginning of the session, female rats had a greater number of post-cluster pauses than their male compatriots drinking the 5% alcohol concentration cocktail (**Fig. 4d**). There were no significant contributions of sex in rats drinking the fructose cocktail at any concentration of alcohol.

When analyzing the number of pauses after an extended run of licks, the overall behavior was similar to the shorter post-cluster pauses (**Fig. 4f-m**). That is, rats demonstrated similar sex and cocktail specific effects that changed during the session, with male rats showing a greater sensitivity to alcohol concentration and cocktail type than female rats. In the beginning of the session, the number of pauses after a run (**Fig. 4f-i**) generally mirrored those seen in post-cluster pauses. A similar interaction between alcohol and cocktail type was seen in male rats (**Fig. 4g**), although there were no alcohol concentration-specific differences at this level of analysis, unlike post-cluster pauses. For the remainder of the session, male rats showed significantly greater numbers of run pauses when drinking lower alcohol concentration glucose cocktails than equivalent fructose cocktails (**Fig. 4k**), while female rats did not (**Fig 4j**).

#### These significant differences in pauses after a cluster of licks and to a lesser degree after a run of licks at the beginning of the session may reflect several different processes

During this part of the session, the direct central effects of the cocktails should be relatively minimal. One explanation for these effects may be that post-ingestive effects like satiety effects are occurring dynamically and early in the session (Davis, 1989; Schier and Spector, 2016). As detectable levels of both consumed glucose and alcohol are present in the brain very soon after the start of drinking, the cocktails may have an impact on brain neuronal firing and behavior much sooner than expected. A second explanation for these behaviors may be that cluster pause number at the beginning of a session better represent processes beyond satiety alone.

Indeed, while post-ingestive control is often framed as negative feedback and satiety (Davis, 1989), post-ingestive feedback, particularly of carbohydrate consumption, can also be positive feedback (Sclafani, 2001). As post-cluster pauses are linked to the number of times a rat returns to drink, this behavior during the early parts of the session may also reflect broader differences in motivation for a particular cocktail, perhaps through associative learning mechanisms (Ostlund et al., 2013; Kosse et al., 2015; Mendez et al., 2015). While direct comparisons of pauses between early and late time intervals within 4-hour long sessions has not been performed previously, other groups have reported on these behavioral measures for sessions up to 2 hours long (Davis and Perez, 1993; Spector and St. John, 1998; Spector et al., 1998).

#### Clearly, additional work will need to be conducted to carefully evaluate the possible biological mechanisms regulating this post-ingestive and satiety-linked behavior

It should also be noted that the number of pauses after a run of licks seems to be more sensitive to sex differences than the shorter cluster pauses. In our study, run-based post-ingestive measures in male rats during the later portion of the session appeared to be more responsive to changes in the concentration of alcohol in the glucose cocktail than the fructose cocktail. In female rats, run-based post-ingestive measures during this part of the session appeared to be overall less sensitive to the changes in the interaction between alcohol concentration and cocktail type.

Run pauses also emphasized sex differences across more time periods during the session than did cluster pauses. The presence of sex differences at both early and late time intervals suggests that post-run pauses are more sensitive to sex differences after glucose and alcohol have entered the brain, in addition to the sex differences during the gradual rise soon after the start of drinking. Together, both microstructure categories suggest that male rats are more responsive to the post-ingestive effects of cocktail composition, which is present no matter how much glucose is in the brain, while female rats show more stable satiety responses between different sugar types.

Considering sex differences within glucose or fructose cocktails, only glucose cocktails showed differences in both cluster and run pauses (**Fig. 4d, i**). Male rats drinking glucose cocktails in the first 30 minutes of the session showed a steady decrease in cluster and run pauses over alcohol concentration for both post-cluster and post-run pauses, suggesting a rise in post-ingestive effects such as satiety. On the other hand, female rats exhibited an inverted U-shaped curve in pause number when alcohol concentration increased. In the context of dose-response curves, both cluster and run pauses in male rats resembles a leftward dose-response shift when compared to female rats. The widely accepted pharmacological interpretation of a leftward shift suggests that male rats are more sensitive to the post-ingestive negative effects of increasing alcohol concentration compared to female rats. This effect is a uniquely glucose-alcohol cocktail effect, as it was not observed in fructose cocktails. In the future, we will need to determine the mechanisms regulating these post-ingestive behaviors. Further, sex as a biological variable will have to be systematically investigated in future studies.

### Timing of post-ingestive effects varies by sex at short pause durations

In addition to the total number of pauses after a cluster or run of licks, the length of the pause itself can also vary. In our study, only the number of pauses shorter than 10s were significantly impacted by the cocktail type (**Fig S3**). The interpretation of pause length remains debated. While some groups have considered pauses greater than 0.5s to signify termination of licking by behaviors other than drinking (Davis and Smith, 1992), others have categorized pauses between 0.5s-1s to be a mechanical consequence of multiple missed licks or mouth movements in response to palatable or aversive tastes (Grill and Norgren, 1978; Spector et al., 1998). In contrast, pause lengths of 1-10s are long enough for other behaviors to interrupt drinking. Some behaviors that commonly occur around bouts of drinking are locomotion, grooming, rearing, and sniffing (Bolles, 1960; Silverman, 1965; Grill and Norgren, 1978; van Dam et al., 2013; Schier and Spector, 2019). These shortest pauses (0.5s-1s) between clusters may reflect different behaviors than 1-10s pauses, which will need to be empirically determined. Moreover, some researchers define longer pauses (i.e. greater than 300s) as markers of the end of a “meal” (Spector et al., 1998). In this context, in our data, longer length pauses (10-300s+) were not impacted by the cocktail type, and therefore cocktail type did not impact the number of “meals” during the session.

When evaluating the interaction between the number of pauses after a cluster or run of licks, the duration of the pause, cocktail type, alcohol percentage, and sex, several trends are notable. First, pauses after a cluster or run of licks both follow similar trends when the alcohol concentration is low (**Fig. 5a-d, i-l**). With low alcohol cocktails, significant differences in pause number only appear in the later portion of the session. Both glucose and alcohol concentrations gradually rise in the brain soon after the start of drinking (Nurmi et al., 1999; Wakabayashi and Kiyatkin, 2015), and alcohol starts to arrive in the brain around the same time. Therefore, the first 30 minutes of the session represents an interval of rising brain glucose and alcohol, while the rest of the session represents an interval after glucose and alcohol have entered the brain and presumably exerted central effects. Additionally, other processes may take effect alongside satiety to impact drinking, such as post-ingestive positive feedback (Sclafani, 2001). In this context, our finding that male rats drinking the glucose cocktail pause more often than rats of both sexes drinking the fructose cocktail suggests that male rats drinking glucose cocktails are either less sensitive to post-ingestive satiety signals, or there is greater positive feedback when drinking glucose cocktails.

As the alcohol concentration increases to 5-10%, there are nuanced interactions among cocktail type, session time, and sex. Mainly, significant differences in pause numbers appear within the first 30 minutes of the session much more often than later in the session. Early in the session, male rats drinking higher alcohol concentration glucose cocktails often pause significantly less often than their equivalents of both sexes drinking the fructose cocktail (**Fig. 5e, g, o**). Within our current conceptual framework, one possible pharmacokinetic explanation of this behavior is that with the higher concentration of alcohol, the centrally mediated mechanisms regulating post-drinking pauses occur sooner within the session (**Fig. 5e-h, m-p**). To answer this, direct measurements of brain glucose and brain alcohol levels would need to be obtained at the same temporal resolution. In addition, at 5% alcohol, some unique sex differences in post-drinking pauses are apparent. Female rats drinking the glucose cocktail at 5% alcohol pause more often for 1-10s after a cluster or run than rats of both sexes drinking the fructose cocktail early in the session (**Fig. 5e, m**). The mechanism for this sex difference at 5% alcohol is not clear, though it should be noted that the sex difference disappears later in the session, as with 10% alcohol cocktails. One possible explanation could be that the post-ingestive effects occur at different rates in male rats than in female rats.

Taken together, overall post-ingestive effects (**Fig. 4**) are a result of an interaction between alcohol, cocktail type, sex, and time within the session influencing the number and duration of pauses. These post-ingestive centrally mediated behaviors may be occurring more rapidly as a function of alcohol concentration and/or sex. Overall, this has important theoretical implications as these post-ingestive effects have been linked to satiety or positive feedback (Davis, 1973; Davis and Smith, 1992; Spector et al., 1998; Sclafani, 2001; Naneix et al., 2020).

### Alcohol intake depends largely on post-ingestive effects

In sum, we found that cocktail type, sex, and time within the session influenced both the volume of cocktail drank and the estimated alcohol dose. The key finding of this study was that the estimated dose of alcohol consumed by the rats depended largely on the post-ingestive effects of each cocktail type. We also found a strong contribution of sex differences to this effect (discussed in further detail below). In comparison to fructose cocktails, glucose cocktail drinking resulted in a leftward shift of the dose-response curve for both sexes **(Fig. 2i-j)**, indicating that added glucose potentiates alcohol drinking compared to added fructose. The primary measure of post-ingestive effects, post-cluster and -run pauses, show the most consistent differences later in the session, and differences in run pauses during this time period are restricted to male rats (**Fig. 4i, r-u**). These differences between glucose and fructose cocktails during this time are restricted to lower alcohol cocktails. Subsequent analyses showed that these differences in pauses were limited to shorter pause durations (0.5-10s; **Fig. S3**). Differences in pause number between cocktail types depended on the alcohol concentration. As alcohol concentration increased, differences in pause number across cocktail type occurred earlier within the session (**Fig. 5**). Therefore, we infer that the dose-dependent differences in estimated alcohol dose drank across the two cocktails are largely mediated by changes in these pauses. In contrast, while palatability is likely a contributing factor in both the volume of cocktail drank and the alcohol dose consumed, palatability in our study appeared to remain constant throughout the session (**Fig. 3**). Assessing the relative contribution of both palatability and post-ingestive effects will require future studies specifically designed to determine these factors.

### Sex differences in post-ingestive effects

As there have been no systematic comparisons of satiety measures across sex with licking microstructures, (Ackroff and Sclafani, 2004), it is currently unknown how satiety in drinking differs across sex. A common conception is that smaller animals (such as female rats) would require a smaller volume of cocktail (**Fig. 2c, d**) to achieve satiety and would show increased effects of satiety (**Fig. 4c-d, l-m**). However, smaller animals also show greater calorie intake per body weight than larger animals because of increases in metabolism (Wang, 1925; Eriksson, 1969), though this may depend on age (Quirós Cognuck et al., 2020). In our study, as female rats consumed equivalent calories as male rats for all cocktails (**Fig. 2g, h**), greater metabolic rates in female rats may account for greater cocktail consumption per body weight.

Namely, the overall increased alcohol intake in female rats may be explained by equivalent (**Fig. 4e**) or lower satiating effects (**Fig. 4a-d**) of each cocktail for female rats despite their lower body weight **(Fig. S2)**. Another explanation for increased intake in female rats could be greater post-ingestive positive feedback, so both satiety and positive post-ingestive feedback will need to be investigated across sex in the future. The interaction of glucose and alcohol in the brain seems to have very different effects on female and male rats. First, the wide differences in alcohol drinking across sex at high alcohol glucose cocktails **(Fig. 2l)** appear to follow sex differences in post-ingestive measures early in the session **(Fig. 4d)**. Second, the timing of post-ingestive effects varies across sex for high-alcohol glucose cocktails (**Fig. 5e-h, m-p**), suggesting that each sex has distinct behavioral responses to the arrival of glucose and alcohol into the brain. In the future, understanding how glucose and alcohol interact in the brain to exert behaviorally relevant effects will need to be considered in studies investigating these sex differences.

### Limitations and future directions

This study reports only naturalistic drinking behavior of sugar-sweetened alcohol cocktails. Our findings support the idea that central regulation is critical for differences seen in cocktail drinking behavior. Several nutrient-sensing systems in the brain have been implicated in both glucose and alcohol drinking (Burdakov et al., 2005; Cippitelli et al., 2010; Karlsson et al., 2016). Future studies will need to be undertaken directly manipulating these neuronal systems to determine their contribution to the differences in behaviors reported here. This hypothesis also predicts that the activity of nutrient-sensing neurons in the brain will vary depending on the type of cocktail being consumed.

In addition to the effects of sugar types on the brain, how sugars affect the body will need to be addressed, particularly considering how fructose accelerates the metabolism of alcohol (Ylikahri et al., 1976; Berman et al., 2003; Villalobos-García et al., 2021) and the distinct metabolic consequences of glucose and fructose (Thorburn et al., 1989).

In our study, we analyzed palatability and post-ingestive measures by varying only the alcohol concentration while holding the sugar concentration constant. As humans often drink glucose and fructose together in the form of high fructose corn syrup, varying the sugar concentration may also provide additional clarity in the interpretation of our data. Additionally, systematically studying drinking with glucose and fructose alone, as well as glucose-fructose mixtures, will help to better understand high fructose corn syrup consumption. All alcohol doses reported in this paper are estimations based on volume drank and body weight; correlating blood alcohol levels with cocktail drinking behavior will be valuable in analyzing how different sugar types influence alcohol metabolism (Czachowski et al., 1999; Villalobos-García et al., 2021). Finally, testing these variables with other models such as chronic alcohol exposure or models of AUD will help to better understand the role of flavored alcoholic beverages (FABs) in AUD.

Additional research will need to be done to parse the contributions of satiety and post-ingestive positive feedback in the number of clusters or runs (here the number of pauses after a cluster or run), especially when considering beverages that contain centrally active components like glucose or alcohol. While the interpretation of different pause lengths (0.5-1s, >1s) remains debated, data derived from precise ethology with microstructure data may further elucidate the true nature of pause length. Current technology allows us to investigate precise ethology mapped to microstructure data, something not possible in previous decades (Grill and Norgren, 1978). Additionally, different groups use varied microstructure criteria (i.e., >0.5 or >1s ILI), which creates difficulty in comparing results across studies. Reporting multiple microstructure lengths may help resolve issues with the interpretation of behavioral data (Naneix et al., 2020). Future studies may also reveal longer microstructures to be behaviorally relevant and facilitate their inclusion in subsequent findings (Lin et al., 2013).

Finally, other physiological variables will need to be studied to understand the impact of FAB drinking. First, while we investigated sex, other biological variables such as age play a critical role in FAB drinking (Griffiths and Sutherland, 1998; Fortunato et al., 2014; Albers et al., 2015). It will also be critical to determine how glucose sensing-neurons regulate cocktail drinking during rising glucose and alcohol levels.

### Translational implications

Our findings highlight the translational importance of sugar type within alcohol drinking, particularly with FABs. FABs vary in their alcohol content but also in the type of sugars added; these sweeteners are thought to influence alcohol intake particularly in adolescents primarily by improving a drink’s taste and palatability (Albers et al., 2015; Sardarian et al., 2020). In our study, glucose added to alcohol shifted the alcohol intake dose response curve, yielding more calories drank and higher alcohol intake than fructose-sweetened alcohol at equivalent concentrations. Further, glucose enhanced measures of palatability during drinking of low alcohol concentrations in a sex- and time-dependent manner. Male rats were especially sensitive to interactions between sugar type and alcohol concentration, whereas female rats showed more stable microstructure across sugar type. Yet females consistently consumed more alcohol relative to body weight and showed unique responses to increasing alcohol concentration, emphasizing sex as a key biological variable in both palatability and post-ingestive measures. These observations are noteworthy given that many commercially available alcoholic beverages contain mixed glucose–fructose sweeteners; the present results suggest that the glucose component facilitates early intake and modifies within-session satiety and/or post-ingestive positive feedback, potentially increasing cumulative alcohol exposure. Our findings support greater disclosure of the composition of sweeteners in product labeling.

Further, our findings support systematic preclinical modeling of added sugars with and without central effects when evaluating the reinforcing efficacy of alcohol across concentrations.

## Data Availability

Data will be made available on request.

## Acknowledgments

This project was made possible by support from the Rural Drug Addiction Research (RDAR) Center (COBRE: P20GM130461) to K.T.W. The authors thank Jessica D. Vollan, Grant Hatcher, and Joshua A. Jolton for assistance with experimental procedures and Dr. Rick Bevins for helpful comments on a draft of this manuscript.

---

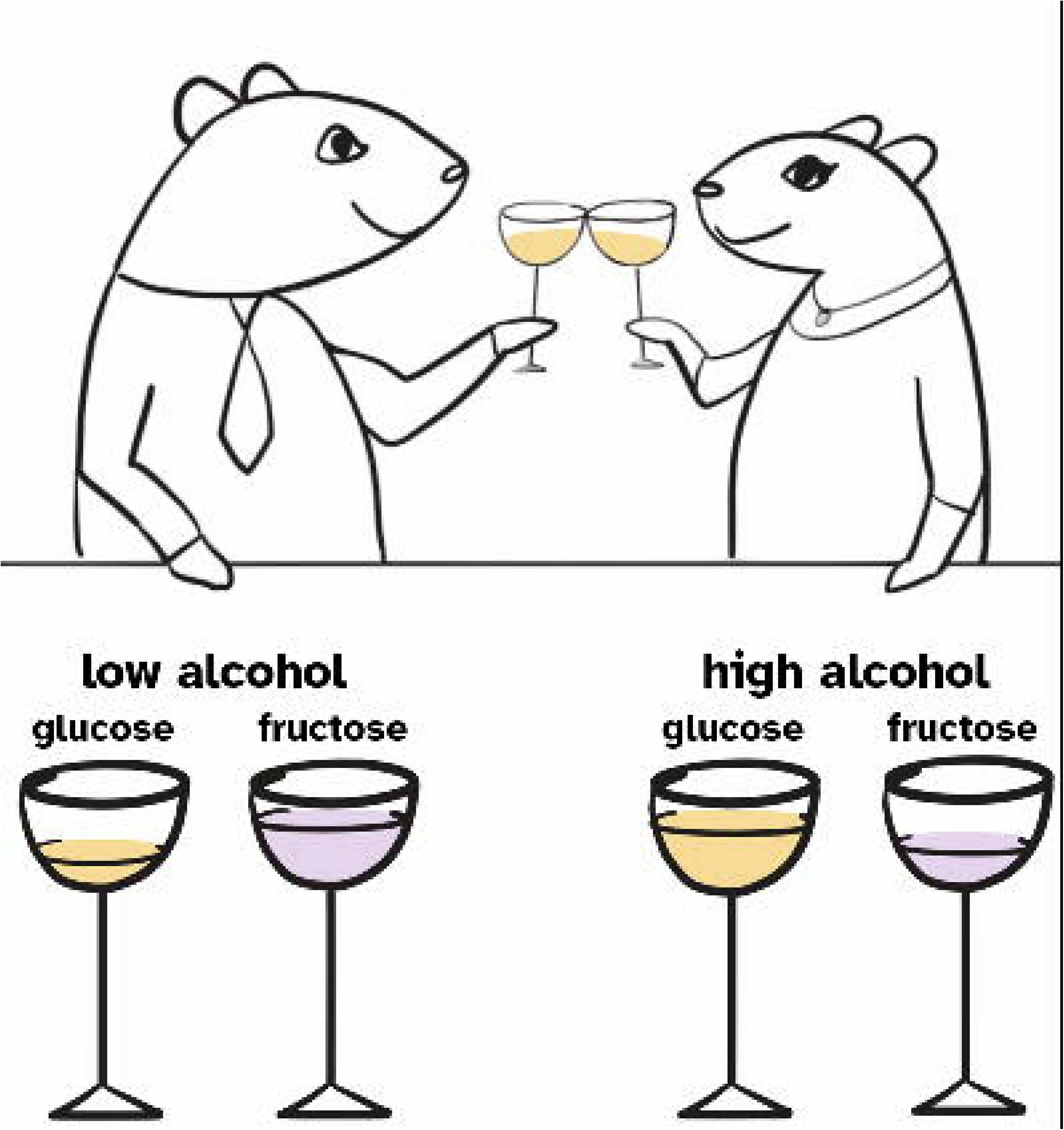

