## Supplemental data for "Glucose and fructose differently mediate alcohol cocktail drinking in female and male rats: interaction of glucose and alcohol on post-ingestive behavior"

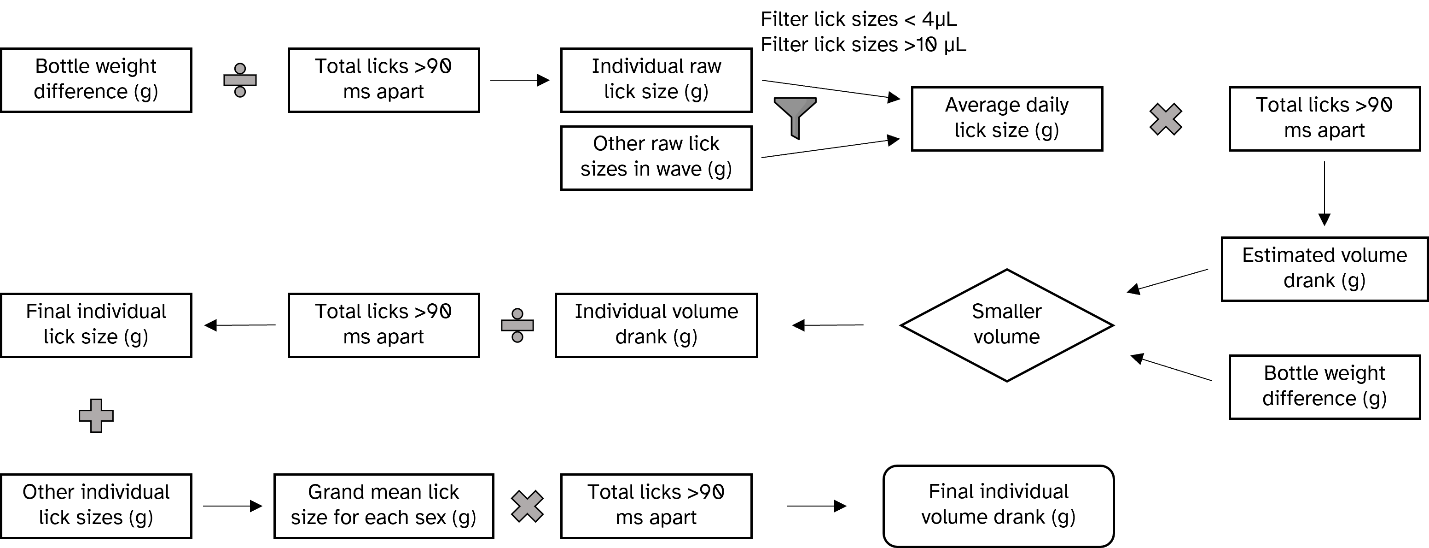

**Fig. S1** **Lick volume calculation workflow**

Daily bottle weights and total lick numbers were used to generate a grand mean lick size for each sex, which was used to calculate the volume drank during each session.

| **Description of analysis** | | **Figure subpanels dependent on this analysis** | **Factors in analysis** | **Fixed effect of alcohol** | **Fixed effect of sugar or pause duration** | **Fixed effect of sex** | **Significant interactions** | **Sex difference** |
| --- | --- | --- | --- | --- | --- | --- | --- | --- |
| Volume drank | entire session | 2a-2d | alcohol, sugar type, sex | *F_3,180_*=205.9, *p*<0.0001 | *F*_1,60_=15.90, *p*=0.0002 | ns | *alcohol dose* x *sugar type F*_3,180_=48.61, *p*<0.0001; *alcohol dose* x *sugar type* x *sex F*_3,180_=4.609*, p=*0.0039 | yes |
| Calories drank | entire session | 2e-2h | alcohol, sugar type, sex | *F*_3,180_=54.92, *p*<0.0001 | *F*_1,60_=7.416, *p*=0.0085 | ns | *alcohol dose* x *sugar type F*_3,180_=52.92, *p*<0.0001; *alcohol dose* x *sugar type* x s*ex* *F*_3,180_=6.980, *p*=0.0002 | yes |
| Alcohol/kg drank | entire session | 2i-2l | alcohol, sugar type, sex | *F*_3,180_=97.15, *p*<0.0001 | ns | *F*_1,60_=22.46, *p*<0.0001 | *alcohol dose* x *sex F*_3,180_=5.576, *p*=0.011; *alcohol dose* x *sugar type F*_3,180_=45.52, *p*<0.0001; *alcohol dose* x *sugar type* x *sex F*_3,180_=2.978, *p*=0.0329 | yes |
| Licks/cluster | first 30 min | 3a | alcohol, sugar type, sex | *F*_3,180_=32.27, *p*<0.0001 | ns | ns | *alcohol dose* x *sugar type F*_3,180_=10.79, *p*<0.0001 | no |
|  | rest of session | 3b | alcohol, sugar type, sex | *F*_3,180_=63.10, *p*<0.0001 | ns | ns | *alcohol dose* x *sugar type* *F*_3,180_=11.33, *p*<0.0001 | no |
| Licks/run | first 30 min | 3c | alcohol, sugar type, sex | *F*_3,180_=38.65, *p*<0.0001 | ns | ns | *alcohol dose* x *sugar type F*_3,180_=13.92, *p*<0.0001 | no |
|  | rest of session | 3d-3g | alcohol, sugar type, sex | *F*_3,180_=58.05, *p*<0.0001 | ns | *F*_1,60_=11.03, *p*=0.0015 | *alcohol dose* x *sex F*_3,180_=5.384, *p*=0.0014; *alcohol dose* x *sugar type F*_3,180_=9.370, *p*<0.0001 | yes |
| Cluster pause number | first 30 min | 4a-4d | alcohol, sugar type, sex | *F*_3,180_=22.93, *p*<0.0001 | ns | *F*_1,60_=5.909, *p=*0.0181 | *alcohol dose* x *sugar type F*_3,180_=6.427, *p*=0.0004*; alcohol dose* x *sex F*_3,180_=3.033, *p*=0.0307; *alcohol dose* x *sex* x *sugar type F*_3,180_=3.583, *p*=0.0150 | yes |
|  | rest of session | 4e | alcohol, sugar type, sex | *F*_3,180_=62.03, *p*<0.0001 | ns | ns | *alcohol dose* x *sugar type F*_3,180_=12.28, *p*<0.0001 | no |
| Run pause number | first 30 min | 4f-4h | alcohol, sugar type, sex | *F*_3,180_=19.45, *p*<0.0001 | ns | *F*_1,60_=5.027, *p*=0.0287 | *alcohol* *dose* x *sugar* *type F*_3,180_=5.618, *p*=0.0011; *alcohol dose* x *sex F*_3,180_=3.818, *p*=0.0110 | yes |
|  | rest of session | 4j-4m | alcohol, sugar type, sex | *F*_3,180_=39.68, *p*<0.0001 | *F*_1,60_=7.849, *p*=0.0068 | ns | *alcohol* *dose* x *sugar* *type* *F*_3,180_=7.616, *p*<0.0001; *alcohol dose* x *sex F*_3,180_=5.751, *p*=0.0009*; alcohol dose* x *sugar type* x *sex F*_3,180_=3.12, *p*=0.0274 | yes |
| Cluster pause number by pause duration | entire session glucose | S3a-d | alcohol, pause duration, sex | *F*_3,90_=45.757, *p*<0.0001 | *F*_3,90_=142.59, *p*<0.0001 | ns | *alcohol dose* x *pause duration* *F*_9,270_=22.880, *p*<0.0001*; alcohol dose* x *sex F*_3,90_=4.017, *p*=0.01; *alcohol dose* x *sex* x *pause duration interaction F*_9,270_=3.647, *p*=0.0003 | yes |
|  | entire session fructose | S3a-d | alcohol, pause duration, sex | *F*_3,90_=14.247, *p*<0.0001 | *F*_3,90_=150.579, *p*<0.0001 | ns | *alcohol dose* x *pause duration* *F*_9,270_=3.986, *p*<0.0001 | no |
| Run pause number by pause duration | entire session glucose | S3e-h | alcohol, pause duration, sex | *F*_3,90_=28.494, *p*<0.0001 | *F*_2,60_=214.61, *p*<0.0001 | ns | *alcohol dose* x *pause duration F*_6,180_=24.029, *p*<0.0001*; alcohol dose* x *sex F*_3,90_=6.141, *p*=0.0008; *alcohol dose* x *sex* x *pause duration F*_6,180_=6.625, *p*<0.0001 | yes |
|  | entire session fructose | S3e-h | alcohol, pause duration, sex | *F*_3,90_=8.638, *p*<0.0001 | *F*_2,60_=197.147, *p*<0.0001 | ns | *alcohol dose* x *pause duration* *F*_6,180_=3.292, *p*=0.004 | no |

**Table S1. 3-way Mixed Effects Analysis results**

| **Description of analysis** | | **Figure subpanel** | **Factors in analysis** | **Fixed effect of alcohol F value** | **p-value** |
| --- | --- | --- | --- | --- | --- |
| Volume drank | female by sugar | 2a | alcohol, sugar type | *F*_3,90_=89.79 | *p*<0.0001 |
|  | male by sugar | 2b | alcohol, sugar type | *F*_3,90_=118.2 | *p*<0.0001 |
|  | fructose by sex | 2c | alcohol, sex | *F*_3,90_=44.63 | *p*<0.0001 |
|  | glucose by sex | 2d | alcohol, sex | *F*_3,90_=170.4 | *p*<0.0001 |
| Calories drank | female by sugar | 2e | alcohol, sugar type | *F*_3,90_=22.91 | *p*<0.0001 |
|  | male by sugar | 2f | alcohol, sugar type | *F*_3,90_=32.74 | *p*<0.0001 |
|  | fructose by sex | 2g | alcohol, sex | *F*_3,90_=8.401 | *p*<0.0001 |
|  | glucose by sex | 2h | alcohol, sex | *F*_3,90_=75.54 | *p*<0.0001 |
| Alcohol/kg drank | female by sugar | 2i | alcohol, sugar type | *F*_3,90_=58.02 | *p*<0.0001 |
|  | male by sugar | 2j | alcohol, sugar type | *F*_3,90_=39.61 | *p*<0.0001 |
|  | fructose by sex | 2k | alcohol, sex | *F*_3,90_=115.90 | *p*<0.0001 |
|  | glucose by sex | 2l | alcohol, sex | *F*_3,90_=35.08 | *p*<0.0001 |
| Licks/cluster | first 30 min combined by sugar | 3a | alcohol, sugar type | *F*_3,186_=33.0 | *p*<0.0001 |
|  | rest of session combined by sugar | 3b | alcohol, sugar type | *F*_3,186_=63.65 | *p*<0.0001 |
| Licks/run | first 30 min combined by sugar | 3c | alcohol, sugar type | *F*_3,186_=39.05 | *p*<0.0001 |
|  | rest of session female by sugar | 3d | alcohol, sugar type | *F*_3,90_=29.79 | *p*<0.0001 |
|  | rest of session male by sugar | 3e | alcohol, sugar type | *F*_3,90_=41.0 | *p*<0.0001 |
|  | rest of session fructose by sex | 3f | alcohol, sex | *F*_3,90_=21.55 | *p*<0.0001 |
|  | rest of session glucose by sex | 3g | alcohol, sex | *F*_3,90_=39.54 | *p*<0.0001 |
| Cluster pause number | first 30 min female by sugar | 4a | alcohol, sugar type | *F*_3_ _,90_=10.28 | *p*<0.0001 |
|  | first 30 min male by sugar | 4b | alcohol, sugar type | *F*_3,90_=17.85 | *p*<0.0001 |
|  | first 30 min fructose by sex | 4c | alcohol, sex | *F*_3,90_=3.072 | *p*=0.0317 |
|  | first 30 min glucose by sex | 4d | alcohol, sex | *F*_3,90_=24.73 | *p*<0.0001 |
|  | rest of session combined by sugar | 4e | alcohol, sugar type | *F*_3,186_=59.98 | *p*<0.0001 |
| Run pause number | first 30 min female by sugar | 4f | alcohol, sugar type | *F*_3,90_=8.387 | *p*<0.0001 |
|  | first 30 min male by sugar | 4g | alcohol, sugar type | *F*_3,90_=19.31 | *p*<0.0001 |
|  | first 30 min fructose by sex | 4h | alcohol, sex | *F*_3,90_=3.491 | *p*=0.0189 |
|  | first 30 min glucose by sex | 4i | alcohol, sex | *F*_3,90_=17.89 | *p*<0.0001 |
|  | rest of session female by sugar | 4j | alcohol, sugar type | *F*_3,90_=7.453 | *p*=0.0002 |
|  | rest of session male by sugar | 4k | alcohol, sugar type | *F*_3,90_=43.14 | *p*<0.0001 |
|  | rest of session fructose by sex | 4l | alcohol, sex | *F*_3,90_=11.18 | *p*<0.0001 |
|  | rest of session glucose by sex | 4m | alcohol, sex | *F*_3,90_=28.61 | *p*<0.0001 |

**Table S2. *2-way Mixed Effects Analysis fixed effect of alcohol results***

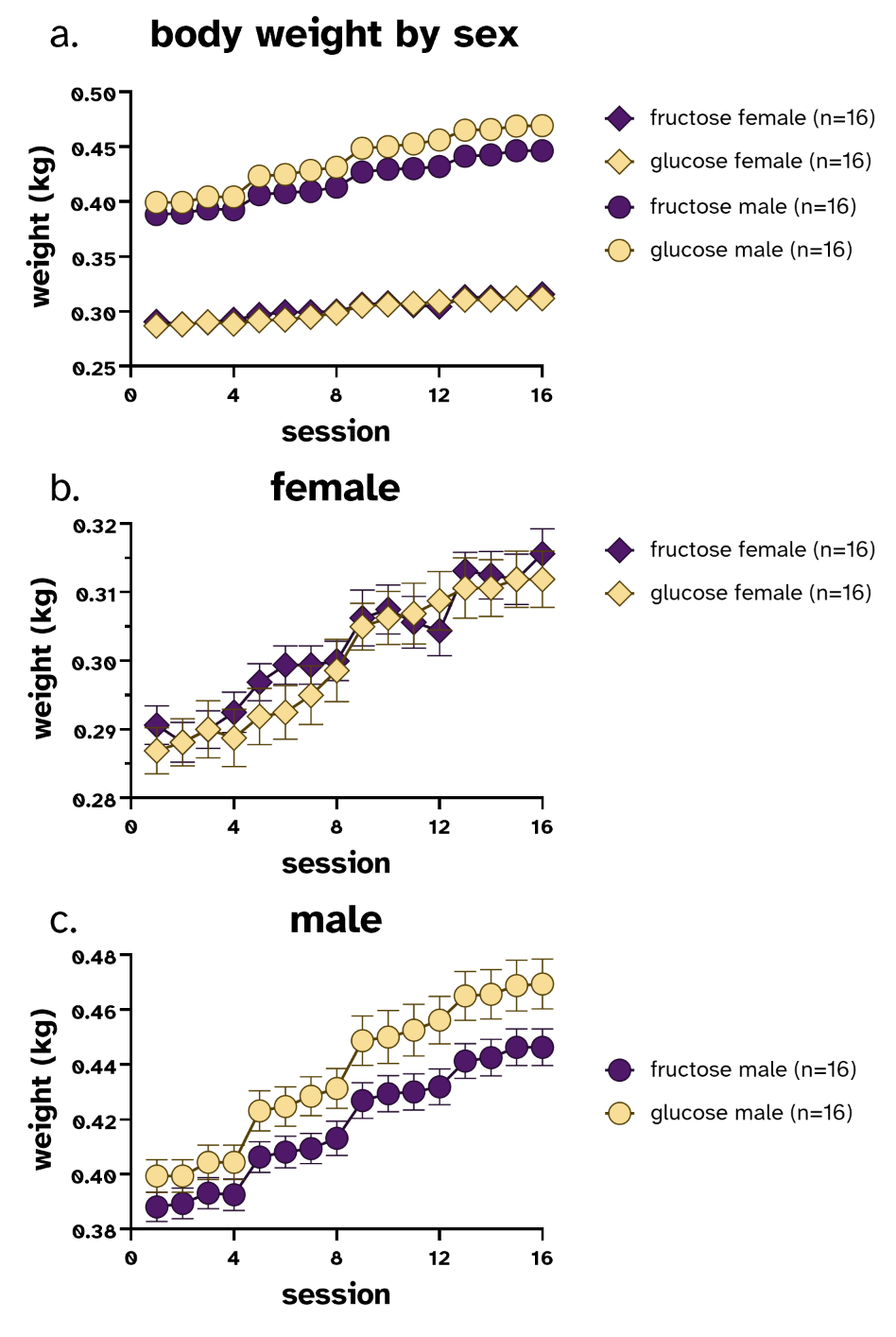

**Fig. S2** **Rat body weights per session. Each set of four sessions represents increasing alcohol concentrations.**

**(a)** Male rats consistently had higher body weights than female rats throughout the 16 sessions. **(b)** Female rats had similar body weights between the two cocktail types at each alcohol concentration, gradually rising in body weight. **(c)** Male rats drinking glucose cocktails increased in body weight more than the rats drinking fructose cocktails.

| **Description of analysis** | **Sex** | **Figure subpanels dependent on this analysis** | **Sugar type** | **Fixed effect of alcohol concentration** | **Fixed effect of pause duration** | **Alcohol concentration x pause duration interaction** |
| --- | --- | --- | --- | --- | --- | --- |
| Cluster pause number by pause duration | female+male combined | S3a-d | fructose | *F*_3,93_=14.618, *p*<0.0001 | *F*_3,93_=148.091, *p*<0.0001 | *F*_9,279_=4.009, *p*<0.0001 |
|  | female | S3a-d | glucose | *F*_3,45_=13.522, *p*<0.0001 | *F*_3,45_=60.15, *p*<0.0001 | *F*_9,135_=7.768, *p*<0.0001 |
|  | male | S3a-d | glucose | *F*_3,45_=37.728, *p*<0.0001 | *F*_3,45_=37.728, *p*<0.0001 | *F*_9,135_=18.838, *p*<0.0001 |
| Run pause number by pause duration | female+male combined | S3e-h | fructose | *F*_3,93_=8.855, *p*<0.0001 | *F*_2,62_=192.436, *p*<0.0001 | *F*_6,186_=3.361, *p*=0.004 |
|  | female | S3e-h | glucose | *F*_3,45_=4.843, *p*=0.005 | *F*_2,30_=73.367, *p*<0.0001 | *F*_6,90_=4.270, *p*=0.0008 |
|  | male | S3e-h | glucose | *F*_3,45_=39.08, *p*<0.0001 | *F*_2,30_=182.834, *p*<0.0001 | *F*_6,90_=32.298, *p*<0.0001 |

**Table S3.** Two-way Mixed Effects Analysis results of pause numbers by pause duration and alcohol concentration, within sex and sugar group

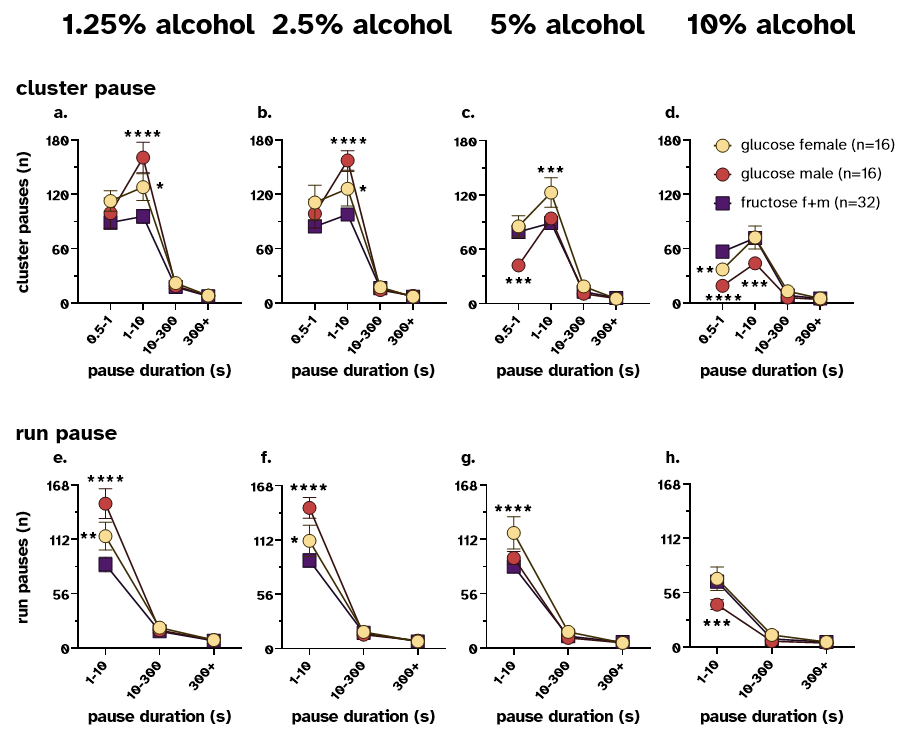

***Figure S3.*** *The number of pauses by pause duration varies by sugar, sex, alcohol concentration, and microstructure length. The most frequent pause length is 1-10s for both cluster and run microstructures. (****a, b****) Rats of both sexes drinking glucose cocktails have significantly more pauses after a cluster that are 1-10s long than rats drinking fructose cocktails at the 1.25% and 2.5% alcohol concentration. (****c****) At 5% alcohol, male rats drinking glucose cocktails had fewer pauses that were 0.5-1s long compared to either female rats drinking the same cocktail, or rats of both sexes drinking the fructose cocktail. Female rats drinking the glucose cocktail had more cluster pauses that were 1-10s long compared to male rats drinking the same cocktail or rats drinking the fructose cocktail. (****d****) At the highest alcohol concentration, rats of both sexes drinking the fructose cocktail had a greater number of pauses that were 0.5-1s long, compared to rats drinking the glucose cocktail. Male rats drinking the glucose cocktail had fewer pauses that were 1-10s long compared to female rats drinking the same cocktail or rats drinking the fructose cocktail. Overall, after a run microstructure, the greatest number of pauses were 1-10s long. (****e, f****) At both 1.25% and 2.5% alcohol, male and female rats drinking the glucose cocktail had a greater number of pauses of this length than rats drinking the fructose cocktail. (****g****) Only female rats had a greater number of 1-10s long pauses after a run microstructure compared to either male rats drinking the same cocktail or rats of both sexes drinking the fructose cocktail. (****h****) Finally, male rats drinking the glucose cocktail had fewer 1-10s length pauses than either female rats drinking the same cocktail or rats drinking the fructose cocktail. *p < 0.05, **p<0.01, ***p<0.001, ****p<0.0001 comparing the female glucose or male glucose group to the female+male fructose group at a given alcohol concentration by Šídák’s multiple comparison test.*

| **Description of analysis** | **Alcohol %** | **Figure subpanel** | **Fixed effect of sugar type** | **Fixed effect of pause duration** | **Sugar type x pause duration interaction** | **0.5-1s** | **1-10s** | **10-300s** | **300+s** |
| --- | --- | --- | --- | --- | --- | --- | --- | --- | --- |
| Cluster pause number by pause duration | 1.25% | S3a | *F*_2,61_=4.034, *p*=0.0226 | *F*_3,183_=192.5, *p*<0.0001 | *F*_6,183_=5.221, *p*<0.0001 | ns | ******** | ns | ns |
|  | 2.50% | S3b | *F*_2,61_=3.582, *p*=0.0338 | *F*_3,183_=169.1, *p*<0.0001 | *F*_6,183_=4.103, *p*=0.0007 | ns | ******** | ns | ns |
|  | 5% | S3c | *F_2_*_,61_=4.009, *p*=0.0231 | *F*_3,183_=181.5, *p*<0.0001 | *F*_6,183_=4.777, *p*=0.0002 | ******* | ******* | ns | ns |
|  | 10% | S3d | *F*_2,61_=6.998, *p*=0.0018 | *F*_3,183_=109.4, *p*<0.0001 | *F*_6,183_=5.556, *p*<0.0001 | ******** | ******* | ns | ns |
| Run pause number by pause duration | 1.25% | S3e | *F*_2,61_=6.771, *p*=0.0022 | *F*_2,122_=247.3, *p*<0.0001 | *F*_4,122_=8.251, *p*<0.0001 |  | ******** | ns | ns |
|  | 2.50% | S3f | *F*_2,61_=5.622, *p=*0.0057 | *F*_2,122_=291.8, *p*<0.0001 | *F*_4,122_=7.459, *p*<0.0001 |  | ******** | ns | ns |
|  | 5% | S3g | ns | *F*_2,122_=258.3, *p*<0.0001 | *F*_4,122_=3.313, *p=*0.0129 |  | ******** | ns | ns |
|  | 10% | S3h | ns | *F*_2,122_=160.9, *p*<0.0001 | *F*_4,122_=3.011, *p=*0.0208 |  | ******* | ns | ns |

**Table S4.** Two-way Mixed Effects Analysis results of pause numbers by pause duration and sex and sugar group, within each alcohol concentration. Columns of each pause duration reflect the most significant comparison between female+male fructose group and male or female glucose groups.
